# Thrombocytopenia Independently Leads to Monocyte Immune Dysfunction

**DOI:** 10.1101/2023.05.10.540214

**Authors:** Chen Li, Sara K. Ture, Benjamin Nieves-Lopez, Sara K. Blick-Nitko, Preeti Maurya, Alison C. Livada, Tyler J. Stahl, Minsoo Kim, Anthony P. Pietropaoli, Craig N. Morrell

## Abstract

In addition to their well-studied hemostatic functions, platelets are immune cells. Platelets circulate at the interface between the vascular wall and leukocytes, and transient platelet-leukocyte complexes are found in both healthy and disease states, positioning platelets to provide physiologic cues of vascular health and injury. Roles for activated platelets in inducing and amplifying immune responses have received an increasing amount of research attention, but our past studies also showed that normal platelet counts are needed in healthy conditions to maintain immune homeostasis. We have now found that thrombocytopenia (a low platelet count) leads to monocyte dysfunction, independent of the cause of thrombocytopenia, in a manner that is dependent on direct platelet-monocyte CD47 interactions that regulate monocyte immunometabolism and gene expression. Compared to monocytes from mice with normal platelet counts, monocytes from thrombocytopenic mice had increased toll-like receptor (TLR) responses, including increased IL-6 production. Furthermore, *ex vivo* co-incubation of resting platelets with platelet naïve bone marrow monocytes, induced monocyte metabolic programming and durable changes in TLR agonist responses. Assay for transposase-accessible chromatin with high-throughput sequencing (ATAC-Seq) on monocytes from thrombocytopenic mice showed persistently open chromatin at LPS response genes and resting platelet interactions with monocytes induced histone methylation in a CD47 dependent manner. Using mouse models of thrombocytopenia and sepsis, normal platelet numbers were needed to limit monocyte immune dysregulation and *IL6* expression in monocytes from human patients with sepsis also inversely correlated with patient platelet counts. Our studies demonstrate that in healthy conditions, resting platelets maintain monocyte immune tolerance by regulating monocyte immunometabolic processes that lead to epigenetic changes in TLR-related genes. This is also the first demonstration of sterile cell interactions that regulate of innate immune-metabolism and monocyte pathogen responses.

## Introduction

Sepsis is a life-threatening syndrome of organ dysfunction caused by dysregulated host responses to a systemic infection (1). Sepsis affects approximately 1.7 million adults each year and contributes to more than 250,000 annual deaths in the United States (2). Dysregulated host responses to infection manifest as an aggravated, uncontrolled, and self-sustaining inflammation which spreads via the circulation. Severe sepsis patients display a profound activation of cytokine networks, including IL-6 and TNFα. IL-6 is associated with a poor prognosis and sepsis severity, and is suggested as a sepsis diagnostic and prognostic biomarker (3–6). Pro-inflammatory cytokines during sepsis prolong and augment the sepsis inflammatory responses and contribute to the development of multiple organ failure (7).

A platelet count below 150,000/μl in humans is clinically defined as thrombocytopenia. The incidence of thrombocytopenia is high in patients with sepsis and thought to be due to infection driven platelet activation and consumption. Many studies have shown an inverse correlation between the degree of thrombocytopenia and sepsis outcomes (8). The level of thrombocytopenia correlates with enhanced cytokine network activation, organ dysfunction, and increased mortality (8–11). For this reason, platelet count is included in ICU severity of sepsis illness scoring systems (12). Although clinical evidence that the platelet count is a predictive biomarker of sepsis severity and hospital mortality is abundant, there is little mechanistic understanding of the link between the severity of thrombocytopenia and sepsis severity.

There is a continually evolving and deepening understanding of platelets in initiating and regulating tissue injury and pathogen responses (13–17). Platelets are now recognized as an integral part of immune activation and regulation (18). Platelets contribute to pathogen killing by direct engulfing and killing bacteria, by releasing bactericidal mediators, and by regulating innate and adaptive immune cell responses and trafficking (19–21). For example, platelets limit bacteria growth and dissemination by activating neutrophil extracellular traps to ensnare bacteria (22, 23). Platelets also influence monocyte polarization and cytokine production in host defenses against infection (24, 25). In experimental sepsis, platelets enhance bacteria clearance and protect against lipopolysaccharide (LPS)-induced septic shock by regulating macrophage functions (26–28). Platelets also regulate immune cell functions and differentiation in healthy conditions as platelet deficient mice and mice lacking platelet factor 4 (PF4) had increased numbers of T helper 17 (Th17) type of CD4^+^ T cells (29), mice lacking platelet derived β2M had more reparative monocytes in basal conditions (30, 31), and thrombocytopenic mice had increased vascular leak (32). These studies suggest that while activated platelets promote an inflammatory immune response to tissue injury or infection, resting platelets have central roles in maintaining immune quiescence in healthy conditions. Platelets may therefore function as an immune rheostat to set the tone of the inflammatory environment.

The immune system has evolved to maintain homeostasis in normal conditions and activate when faced with pathogen challenges or sterile tissue injury. This requires the integration of many immune cells, including platelets and monocytes. Circulating monocytes serve as a first line of host defense to recognize and clear pathogens (33, 34). Monocytes release inflammatory cytokines including IL-6 and TNFα that trigger a robust inflammatory reaction. However, this process needs to be tightly regulated. In sepsis, dysregulated monocyte responses contribute to a “cytokine storm” that leads to exaggerated inflammation, tissue damage and potentially mortality (34). The cellular and molecular mechanisms that regulate monocyte immune phenotypes are complex and include roles for platelets. Monocytes and macrophages can also adapt to prior stimulation and respond differently toward a second challenge with the same or different agonist stimulation. The secondary response can be altered in such a way that monocytes respond more to enhance immune responses (trained immunity) or less strongly to prevent excess immune responses (innate tolerance) (35). Different primary stimuli may induce different training programs. The bulk of evidence suggests metabolic reprogramming and epigenetic remodeling are underlying mechanisms that mediate innate immune training (36). However, endogenous cell interactions regulating innate immune training has not been demonstrated.

Given the abundant evidence indicating that platelets are essential players in the immune system in both health and disease, we hypothesize that circulating resting platelets maintain monocyte immune homeostasis in healthy conditions, and thrombocytopenia may therefore independently lead to immune dysregulation, exacerbating systemic sepsis related inflammatory processes. We have now investigated this using thrombocytopenic mouse models and found that circulating platelets directly maintain monocyte immune homeostasis by programming a monocyte innate immune tolerant phenotype in a metabolic regulatory manner. Using mouse sepsis models, we found that thrombocytopenia resulted in exacerbated inflammatory responses and immune dysregulation. We further demonstrate platelet-monocyte CD47 homotypic interactions suppress monocyte inflammatory responses to toll-like receptor (TLR) agonists through histone methylation and chromatin remodeling. Platelet dependent monocyte tolerance is mediated by an AKT/mTOR induced glycolysis pathway that is critical for monocyte immune programming by resting platelets, the first demonstration of endogenous cell based innate immune programming.

## Results

Monocytes respond to pathogens and tissue injury, as well as maintaining tissue homeostasis. Monocyte phenotypes need to be tightly regulated as changes in monocyte activity, differentiation, and function can lead to disease, including autoimmunity and cardiovascular disease (37). To begin to determine whether circulating platelets participate in maintaining monocyte immune homeostasis, we isolated both bone marrow (BM) and circulating monocytes from wild-type (WT) C57BL/6 mice and thrombocytopenic thrombopoietin (TPO) receptor knockout mice (TPOR*^-/-^*). BM monocytes showed similar expression of *Il6* and *Tnf* between WT and TPOR^-/-^ mice. However, circulating monocytes from TPOR^-/-^ mice had significantly higher *Il6* compared to WT mice (**Fig 1A**). When monocytes were LPS stimulated *ex vivo*, isolated circulating monocytes from thrombocytopenic TPOR^-/-^ mice produced more IL-6 than monocytes from WT mice (**Fig 1B**). This indicated that platelets may limit circulating monocyte *Il6* expression and IL-6 production upon stimulation in normal conditions. Because TPOR^-/-^ mice have potential defects in hematopoietic stem cell production and function (38, 39), we used PF4-Cre × diphtheria toxin receptor(DTR)^fl/fl^ (PF4-DTR) mice as a complementary approach to explore the role of acute thrombocytopenia in regulating monocyte LPS responses (40). PF4-DTR mice express the diphtheria toxin receptor (DTR) in PF4 positive cells (megakaryocytes and platelets)(41) and upon DT exposure, these cells undergo apoptosis leading to a decline in platelet count (40). WT and PF4-DTR mice were injected with DT intraperitoneal (i.p.) on d0. The platelet count in DT treated PF4-DTR mice started to decrease by d1 and reached a nadir at d5, with a rebound by d7 (**Fig 1C**). DT did not induce changes in other blood cell numbers (**Supplemental Fig 1A**). Monocytes circulate for about 2d in mice, so by d3 all circulating monocytes are new post-DT, and by d5 represent at least 2 generations of monocytes following thrombocytopenia. On both d3 and d5 post-DT, plasma from WT and PF4-DTR had similar HMGB1 and S100A8/A9 concentrations, indicating that DT induced thrombocytopenia does not lead to plasma DAMPs that may affect new monocyte functions at these time points (**Supplemental Fig 1B**). We isolated both BM and circulating monocytes from DT treated WT and PF4-DTR mice 5d post-DT, when all monocytes were platelet naïve and there was still a low platelet count. BM monocytes had reduced *Il6* and *Tnf* expression in thrombocytopenic mice, however, circulating monocytes from platelet depleted mice had increased *Il6* expression and secreted more IL-6 in response to LPS *ex vivo* (**Fig 1D-E**), indicating that platelet mediated monocyte immune tolerance is dependent on cell interactions in the peripheral circulation. To further demonstrate this, we isolated monocytes from WT and PF4-DTR mice on d10 post-DT when platelet counts were restored. WT and prior thrombocytopenic mice BM and circulating monocytes had similar *Il6* and *Tnf* expression and cytokine secretion following platelet restoration (**Fig 1F-G**). Together, these data suggest that circulating platelets limit monocyte LPS responses in healthy conditions.

**Figure 1.**
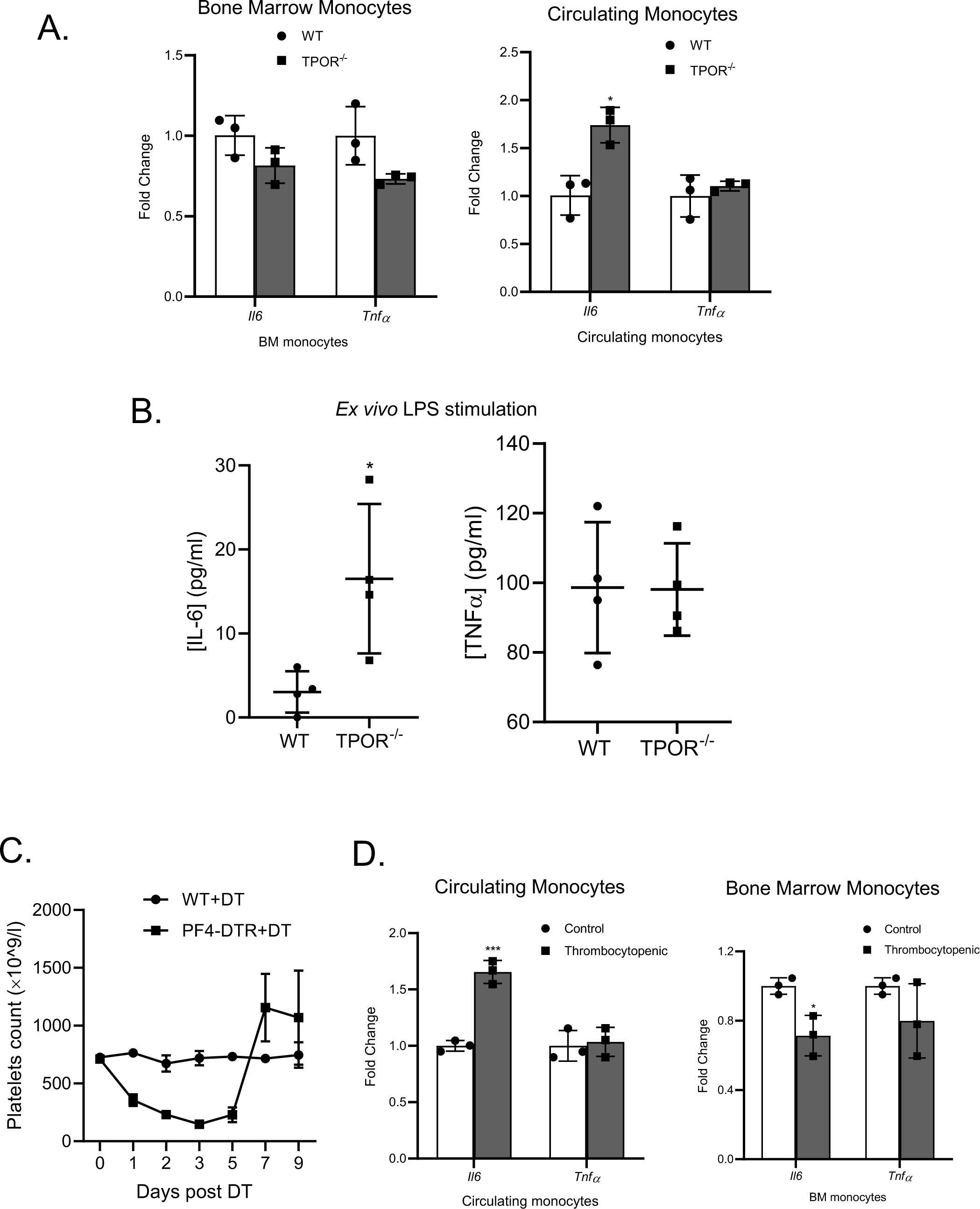
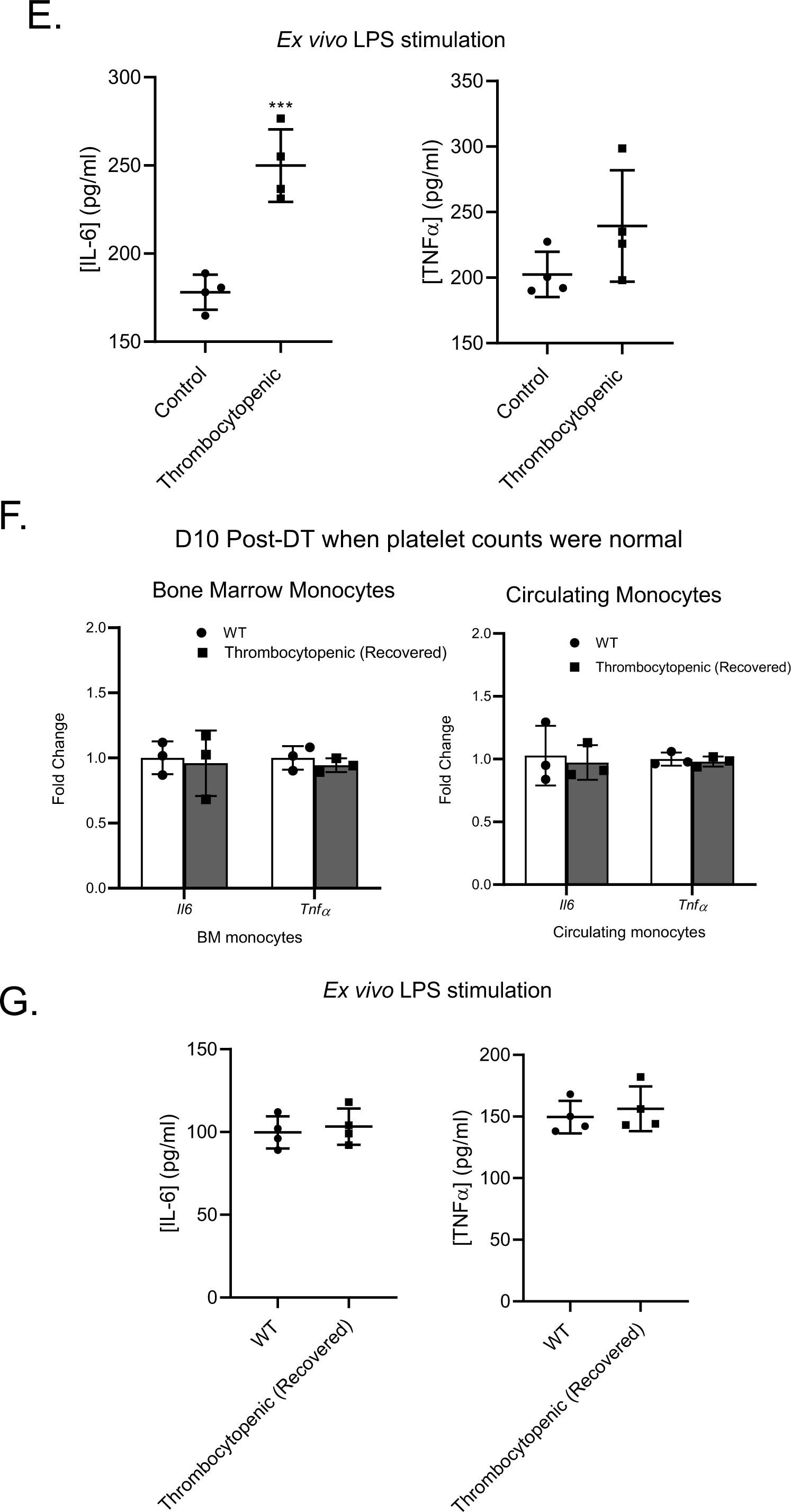
Thrombocytopenia increases circulating monocyte LPS responses. A) Circulating, but not bone marrow (BM) monocytes, from platelet deficient TPOR^-/-^ mice expressed more *Il6* and B) produced more IL-6 *ex vivo* in response to LPS. Isolated circulating monocytes were treated with LPS (50 ng/ml) and 24 hrs later IL-6 and TNFα were determined by ELISA. C) Acute model of thrombocytopenia. PF4^cre^-DTR^flox^ mice or WT control mice were treated with diphtheria toxin (DT) (400 ng/mouse) and platelet counts determined on multiple days post-DT. DT acutely reduced platelet counts for at least 5d. D) Circulating, but not bone marrow, monocytes from acutely thrombocytopenic mice expressed more *Il6* and E) produced more IL-6 *ex vivo* upon LPS stimulation. Isolated circulating monocytes were treated with LPS (50 ng/ml) and 24 hrs later IL-6 and TNFα were determined by ELISA. F) Circulating and BM monocytes from d10 post-DT, platelet count recovered PF4-DTR mice showed similar *Il6* and *Tnf* expression as control mice and G) produced similar IL-6 and TNFα *ex vivo* in response to LPS. Isolated circulating monocytes were treated with LPS (50 ng/ml) and 24 hrs later IL-6 and TNFα were determined by ELISA. Data shown as mean ± SEM. \**P* < 0.05, \*\*\**P <* 0.001, by unpaired, 2-tailed Student’s *t* test.

To determine the *in vivo* functional outcomes of thrombocytopenia on monocyte LPS responses, on d5 post-DT, WT and thrombocytopenic mice were given LPS (5 mg/kg) i.p. to induce sepsis-like responses. 4 hrs post-LPS, plasma IL-6 and TNFα were increased more in thrombocytopenic mice compared to control mice (TNFα likely secondary to the increased IL-6), indicating exacerbated inflammation in the face of thrombocytopenia (**Fig 2A**). Similar results were found using a cecal slurry (CS)-induced polymicrobial sepsis model, in which both DT-induced thrombocytopenic mice (**Fig 2B**) and TPOR^-/-^ mice (**Supplemental Fig 2A**) had increased plasma IL-6 and TNFα 4 hrs after CS was given i.p. Plasma IL-6 and TNFα were similarly increased in thrombocytopenic male and female mice after CS challenge (**Fig 2C**). To determine whether exaggerated inflammatory responses were due to monocytes, we depleted monocytes in both control and thrombocytopenic mice using liposomal clodronate prior to LPS challenge (monocytes were depleted >80%, **Supplemental Fig 2B-C**). Increased plasma IL-6 and TNFα in thrombocytopenic mice were blunted when monocytes were depleted prior to LPS (**Fig 2D**) indicating monocytes are major cell mediators of increased IL-6 in thrombocytopenia. To investigate whether thrombocytopenia also correlated with monocyte dysfunction in human sepsis, we isolated circulating monocytes from confirmed sepsis patient blood at the time of diagnosis, and monocyte *IL6* expression was determined by qRT-PCR and correlated with the platelet counts at the time of blood collection (**Supplemental Table 1**). Monocyte *IL6* expression negatively correlated with patient platelet counts (**Fig 2E**). These data suggest that a decline in platelet count leads to increased propensity for monocyte dysfunction and increased inflammatory cytokines in response to LPS and polymicrobial sepsis.

**Figure 2.**
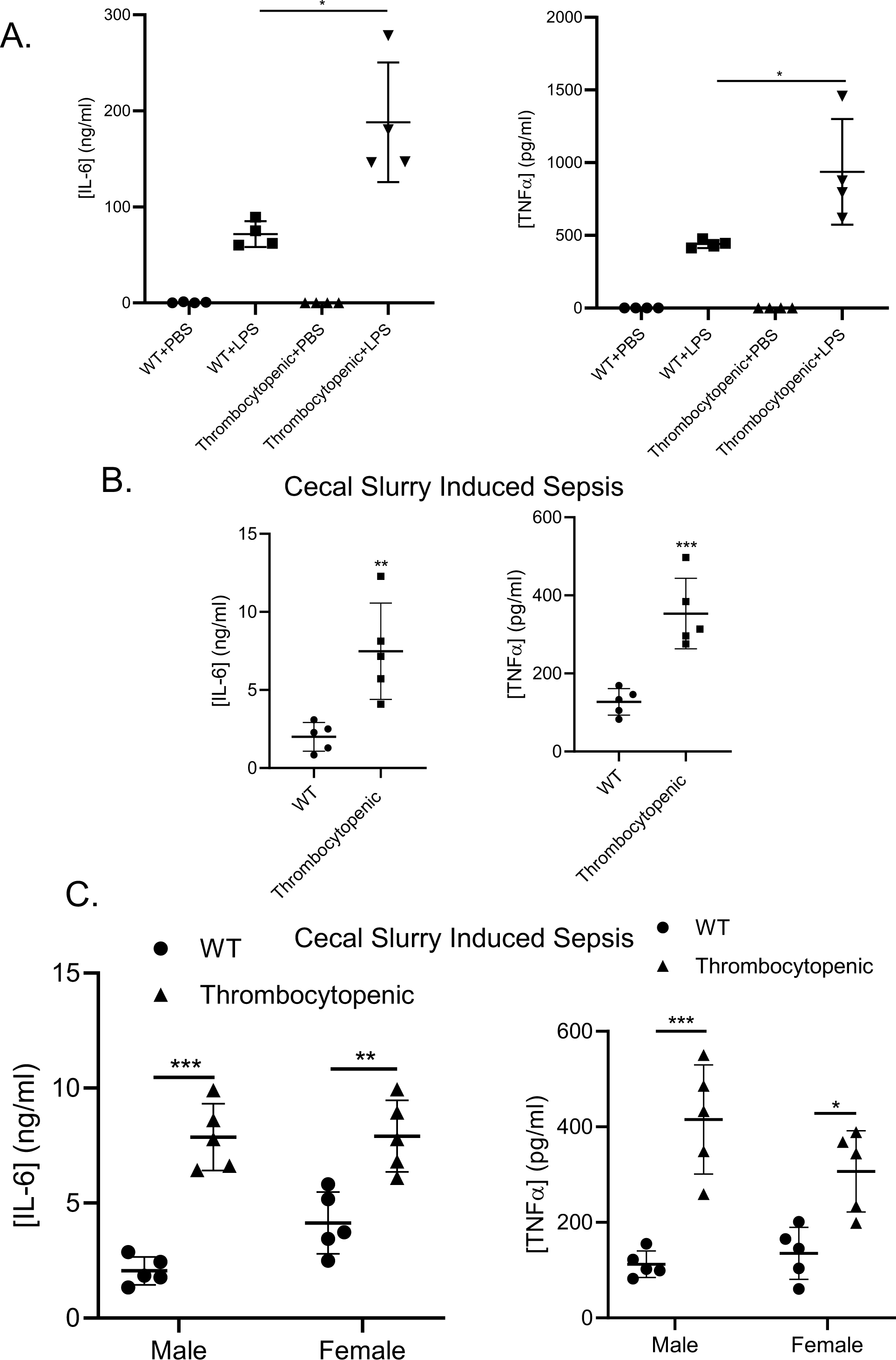
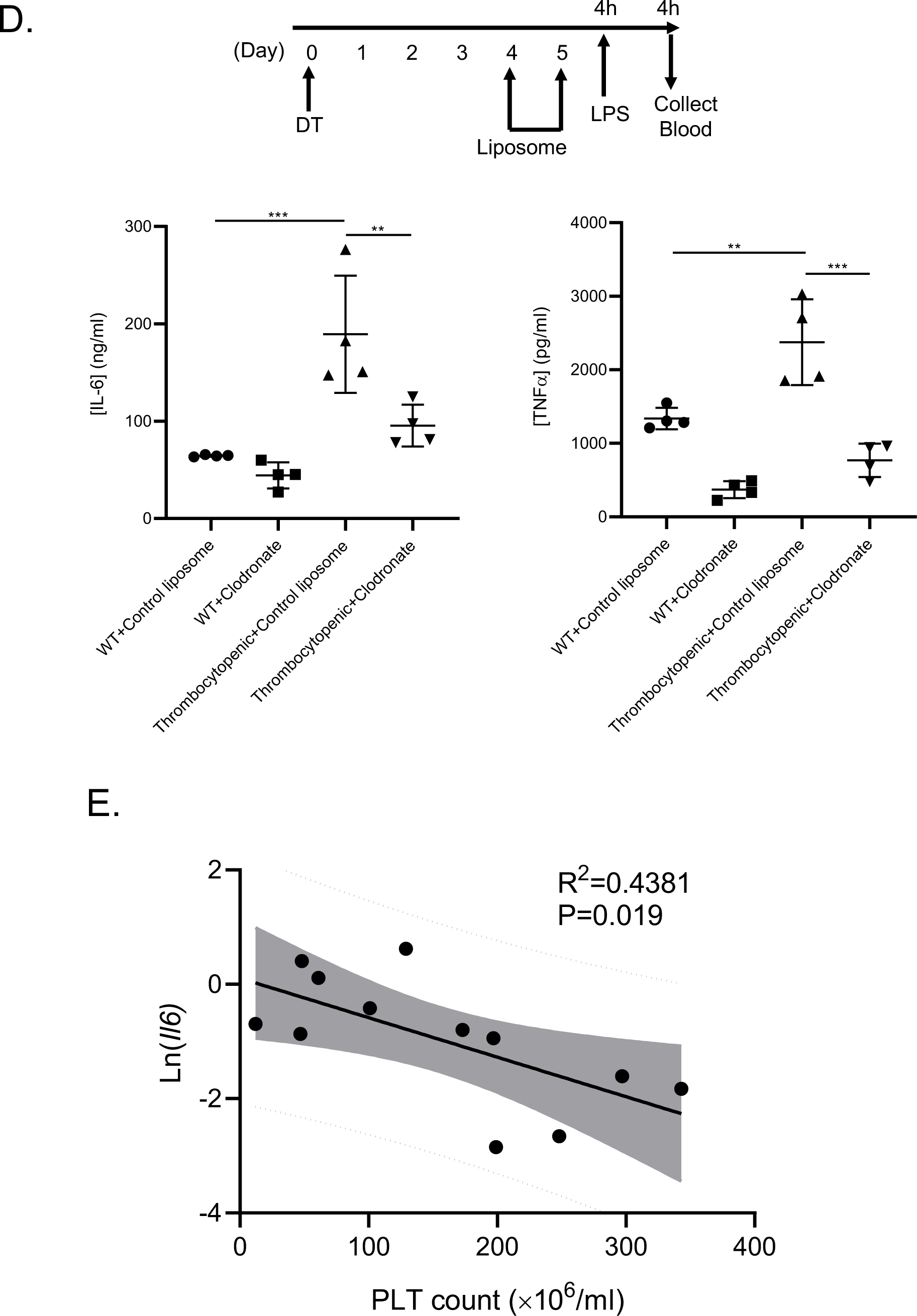
Bacteria associated monocyte signaling is increased by thrombocytopenia *in vivo*. A) Thrombocytopenic mice had increased LPS responses *in vivo*. Mice were treated with LPS (5 mg/kg) intraperitoneally on d5 post-DT and 4 hrs later plasma IL-6 and TNFα were determined by ELISA. (*P < 0.05, by 1-way ANOVA followed by Tukey’s post hoc test). Thrombocytopenic mice had greater IL-6 and TNFα. B) Thrombocytopenic mice had exaggerated responses to polymicrobial sepsis challenge *in vivo.* Control and thrombocytopenic mice were treated with CS (0.8 mg/g) on d5 post-DT and plasma IL-6 and TNFα were determined 4 hrs. (**P < 0.01, ***P < 0.0001, by unpaired, 2-tailed Student’s *t* test). C) Males and females have similar thrombocytopenia associated changes in CS challenge *in vivo.* Age matched male and female WT and thrombocytopenic mice were treated with CS on d5 post-DT and plasma IL-6 and TNFα were determined 4 hrs later by ELISA. (*P < 0.05, **P < 0.01, ***P < 0.0001, by 2-way ANOVA followed by Tukey’s post hoc test). D) Monocyte depletion limited thrombocytopenia associated increase in LPS responses *in vivo*. Monocytes/macrophages were depleted with chlodronate liposomes on d4 and d5 post-DT and 4 hrs after the second chlodronate dose, mice were LPS treated. (**P < 0.01, ***P < 0.0001, by 1-way ANOVA followed by Tukey’s post hoc test). E) Monocyte *IL6* inversely correlated with platelet counts in human sepsis patients. Monocytes were isolated from the peripheral blood of confirmed sepsis patients and mRNA isolated. *IL6* relative to the platelet count was determined by linear regression followed by correlation analysis. Data shown as mean ± SEM.

To mechanistically explore our *in vivo* findings of platelet mediated monocyte LPS tolerance, we performed *in vitro* monocyte-platelet co-culture experiments. Platelet naïve mouse BM monocytes were incubated with resting platelets or control media for 24 hrs at multiple platelet to monocyte ratios. Platelets were then removed by gentle washing and monocytes stimulated with LPS. After overnight co-culture about 90% of monocytes had platelet aggregates, but after gentle washing at the time of LPS stimulation, fewer than 10% of monocytes had platelets still attached (**Supplemental Fig 3A-B**). Pre-incubation of monocytes with platelets reduced IL-6 and TNFα production in a platelet to monocyte ratio dependent manner (**Fig 3A**). Similar results were found using mouse bone marrow derived macrophages (BMDMs) (**Fig 3A**). Both human monocyte cell line THP1 cells and human CD14^+^CD16^-^ circulating monocytes also had less LPS induced IL-6 and TNFα after overnight co-culture with human platelets (**Fig 3A** **and Supplemental Fig 3C**). Despite an attenuation of cytokine production, pre-incubation with platelets increased monocyte and BMDM LPS induced proliferation, as indicated by cell staining dye (CFSE) dilution (**Fig 3B**). This suggests that resting platelet interactions with monocytes may suppresses some cytokine responses, but promote monocyte proliferation, indicating a platelet induced alteration, rather than general suppression, of monocyte LPS responses.

**Figure 3.**
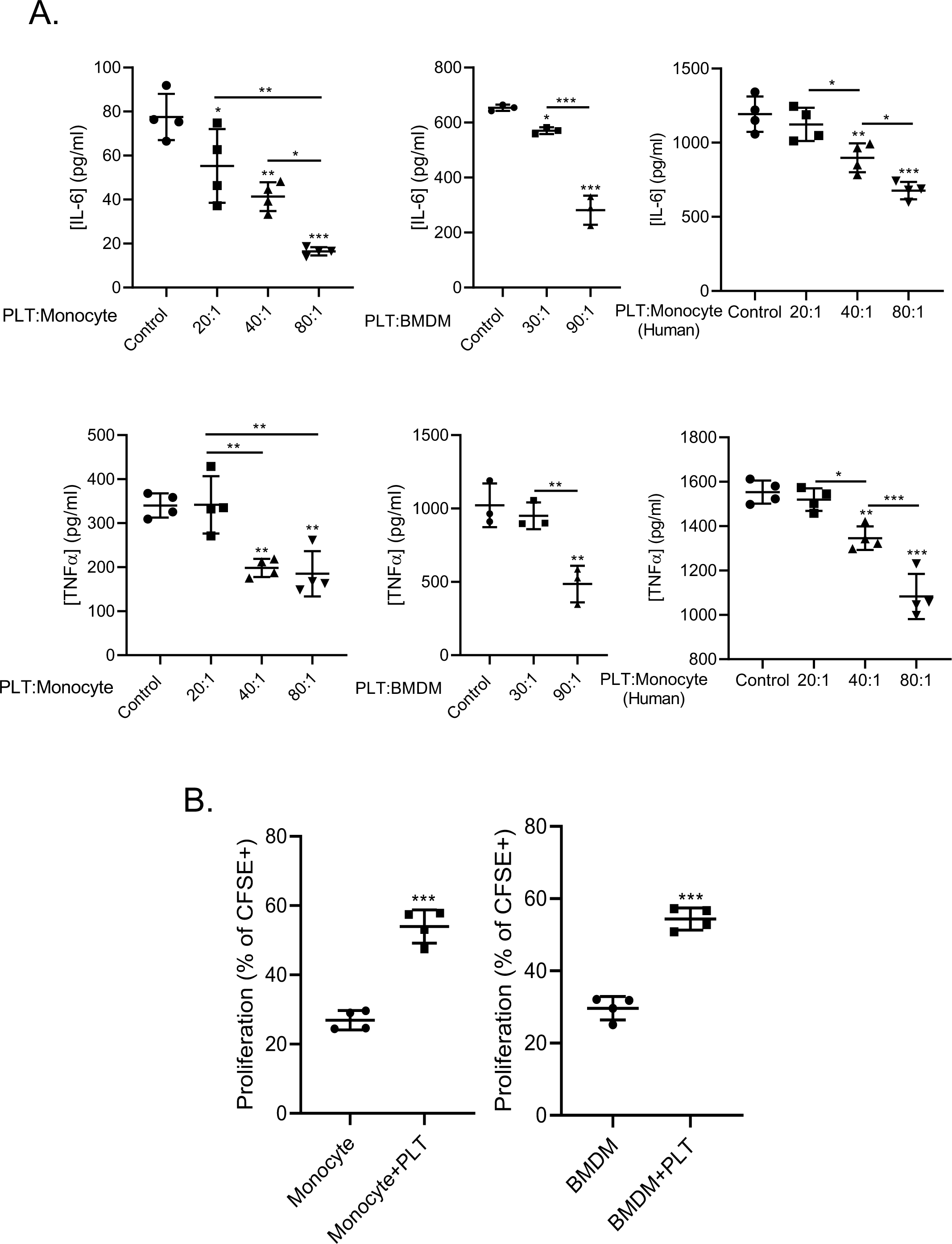
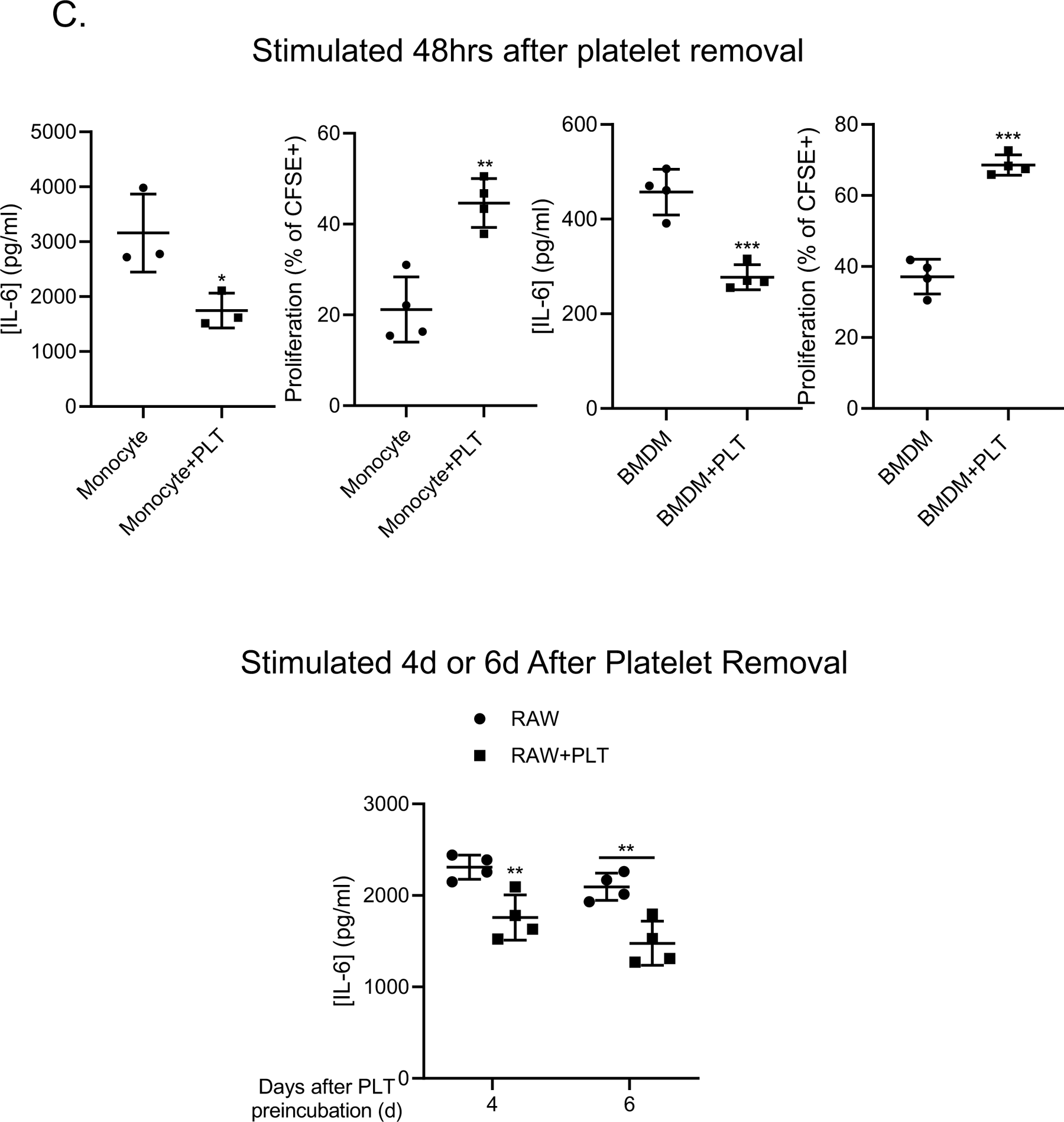
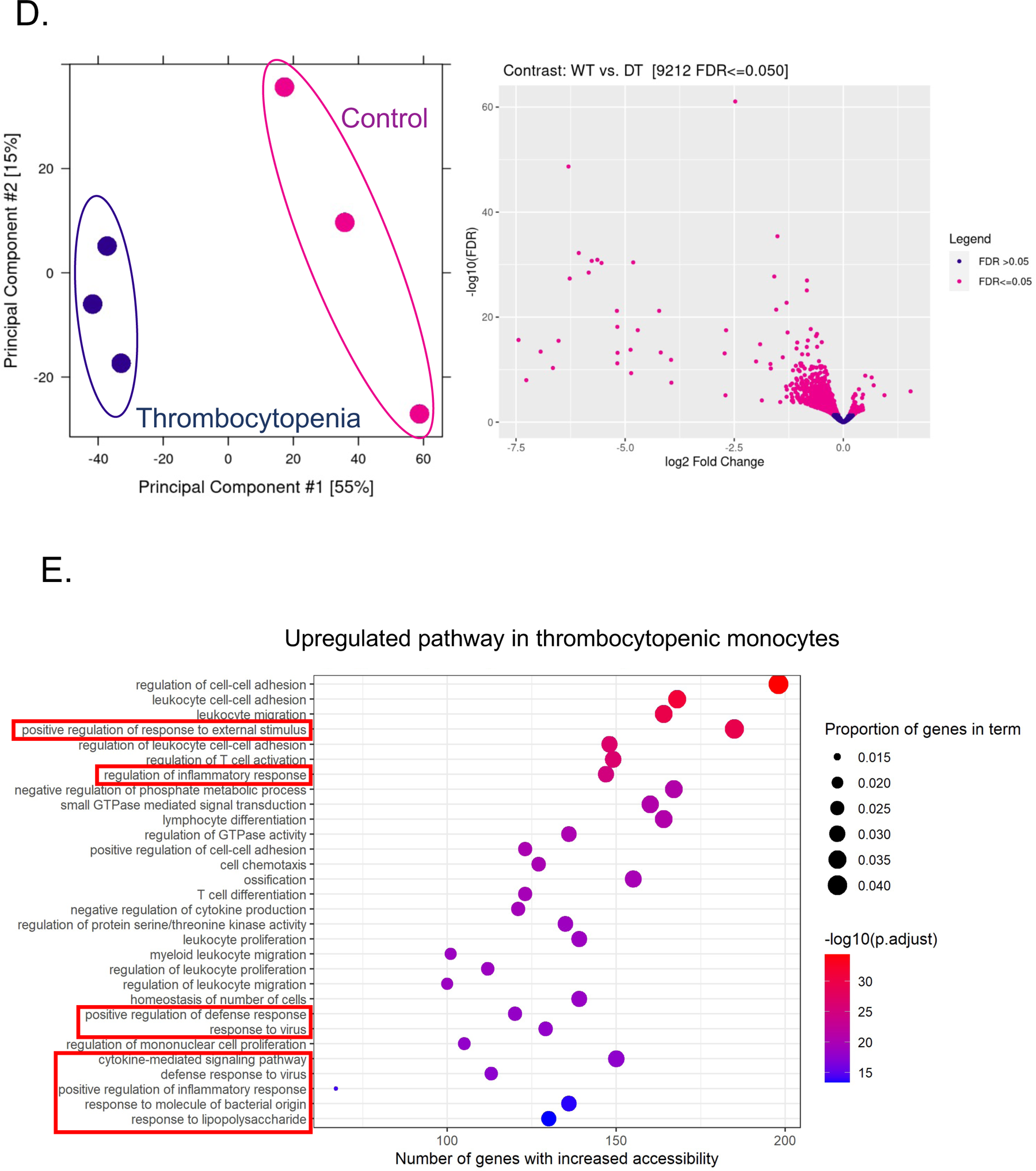
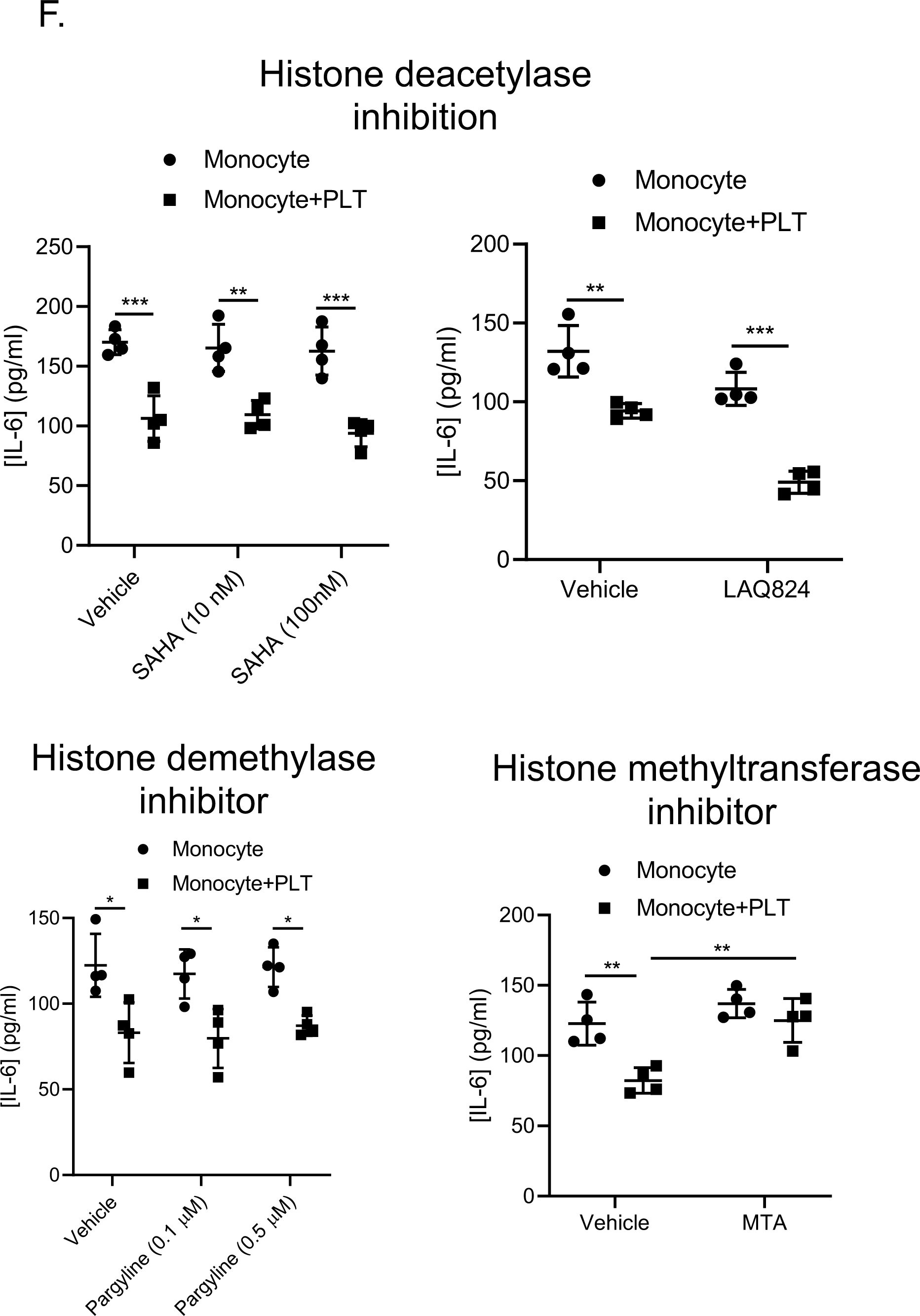
Resting platelets directly mediate monocyte LPS tolerance in a durable manner. Platelets directly mediated monocyte and macrophage LPS tolerance *in vitro*. Primary mouse monocytes, bone marrow derived macrophages (BMDMs) or human CD14^+^CD16^-^ monocytes were co-incubated overnight with platelets and cells then washed to remove platelets. Mono/macs were LPS stimulated and IL-6 and TNFα quantified. (*P < 0.05, **P < 0.01, ***P < 0.0001, by 1-way ANOVA followed by Tukey’s post hoc test). B) Pre-incubation of platelets with monocytes or macrophages increased LPS induced proliferation. Monocytes were CFSE labeled and incubated with platelets overnight prior to washing and LPS stimulation. Despite decreased cytokine responses, prior platelet co-incubation increased monocyte proliferation. (***P < 0.0001 by unpaired, 2-tailed Student’s *t* test). C) Platelet mediated mono/mac immune tolerance is durable for many days after platelet exposure. Platelet naïve monocytes or BMDMs were incubated with platelets overnight and washed to remove platelets. Cells were then LPS stimulated 48 hrs later. Platelet co-incubation limited IL-6 and increased proliferation when stimulated 48 hrs later. Similarly, platelets limited RAW cell LPS induced IL-6 up to 6d after platelet removal. (*P < 0.05, **P < 0.01, ***P < 0.0001 by unpaired, 2-tailed Student’s *t* test). D) Platelets regulate monocyte chromatin remodeling *in vivo*. Circulating monocytes were isolated on d5 post-DT from control and thrombocytopenic mice for ATAC-Seq. Monocytes from thrombocytopenic mice had increased open chromatin patterns. E) Monocytes from thrombocytopenic mice had increased chromatin accessibility in regions related to immune functions. F) Monocytes were incubated with either media or platelets in the absence or presence of histone deacetylase inhibitor SAHA or L4, histone demethylase inhibitor pargyline and histone methyltransferase inhibitor MTA and stimulated with LPS. Data shown as mean ± SEM.

Innate immune cells, including monocytes and macrophages, can durably adapt to previous stimuli and mount greater (trained immunity) or suppressed responses (innate immune tolerance) to secondary stimulation with the same or a different agonist (35, 36, 42–45). To determine whether platelets durably alter monocyte and macrophage LPS responses, we washed away platelets (50:1 ratio) after co-culture and left monocytes or BMDM in culture media for another 48 hrs before LPS stimulation. Monocytes and BMDMs incubated with platelets 48 hrs prior maintained decreased LPS induced IL-6, but had increased proliferation compared to cells incubated with control media (**Fig 3C**). Similar results were found using RAW cells (mouse macrophage cell line) with LPS stimulation up to 6d after platelet removal (**Fig 3C**). These findings indicate that platelet mediated monocyte immune tolerance is durable and therefore likely dependent on gene reprogramming.

Epigenetic reprogramming and changes in chromatin organization are the major molecular basis of innate immune training and tolerance(35). Trained immunity is contingent upon inflammatory gene regions having more open chromatin, and immune tolerance dependent on these regions having more condensed chromatin (42, 46). We therefore hypothesized that circulating monocytes in thrombocytopenic conditions may exhibit more open chromatin at gene loci related to immune responses compared to monocytes from control mice. We collected circulating monocytes from control WT mice and acutely thrombocytopenic mice on d5 post-DT and performed ATAC-Seq to measure chromatin accessibility across the genome. Principal component analysis demonstrated that monocytes from thrombocytopenic mice had a distinct chromatin configuration compared to WT monocytes, with differential accessible chromatin peaks indicating more open chromatin regions (**Fig 3D**). Gene ontology (GO) analysis on regulatory regions that were more accessible in monocytes from thrombocytopenic mice indicated that more accessible chromatin regions were associated with genes involved in the regulation of cell adhesion, inflammatory responses, positive regulation of defense responses, or responses to LPS/TLR related pathways (**Fig 3E**). Altered chromatin accessibility prompted us to investigate whether epigenetic regulators are associated with resting platelet induced monocyte tolerance. Inhibition of histone deacetylases (SAHA, L4) or histone demethylases (pargyline) by specific inhibitors had no effect on monocyte tolerance. In contrast, inhibition of histone methyltransferases with MTA inhibited platelet mediated monocyte tolerant responses (**Fig 3F**), supporting the hypothesis that histone methylation is likely to be involved in the reprogramming of monocytes. Histone H3K9me3 and H3K27me3 are two histone methylation modifications associated with the epigenetic repression of immune-related genes (47, 48). Western blot analysis showed that both H3K9me3 and H3K27me3 increased in platelet co-incubated monocytes compared to control media treated monocytes (**Fig 4C**). In contrast, the level of histone H3K9 acetylation (H3K9ac) was similar in each condition (**Fig 4C**). Increased H3K9me3 and H3K27me3 were also observed in circulating monocytes from WT mice compared to thrombocytopenic mice, indicating platelet mediated monocyte epigenetic modifications *in vivo* (**Fig 4G**). Together, these data suggest that platelet mediated monocyte immune tolerance is durable and associated with histone methylation and chromatin remodeling.

**Figure 4.**
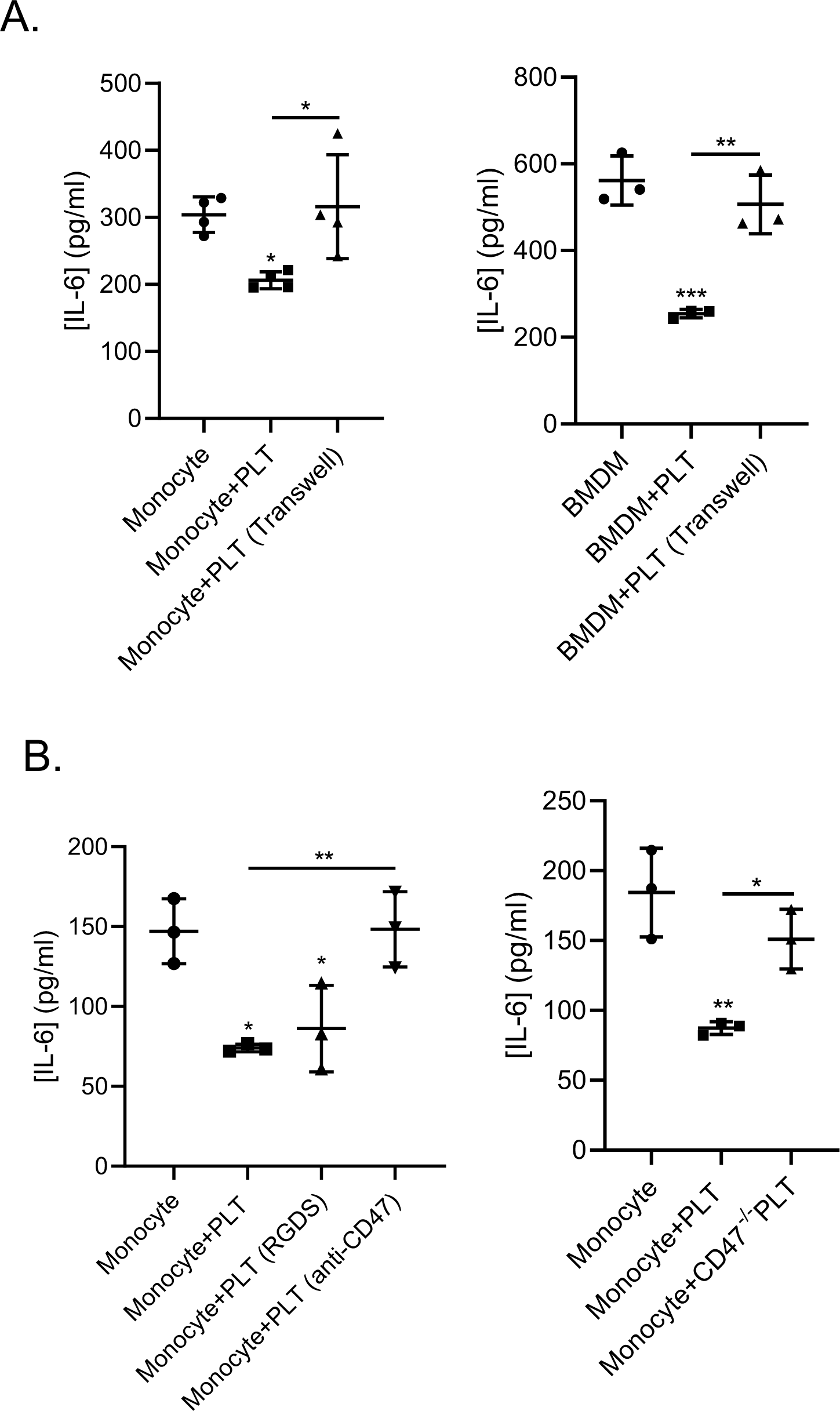
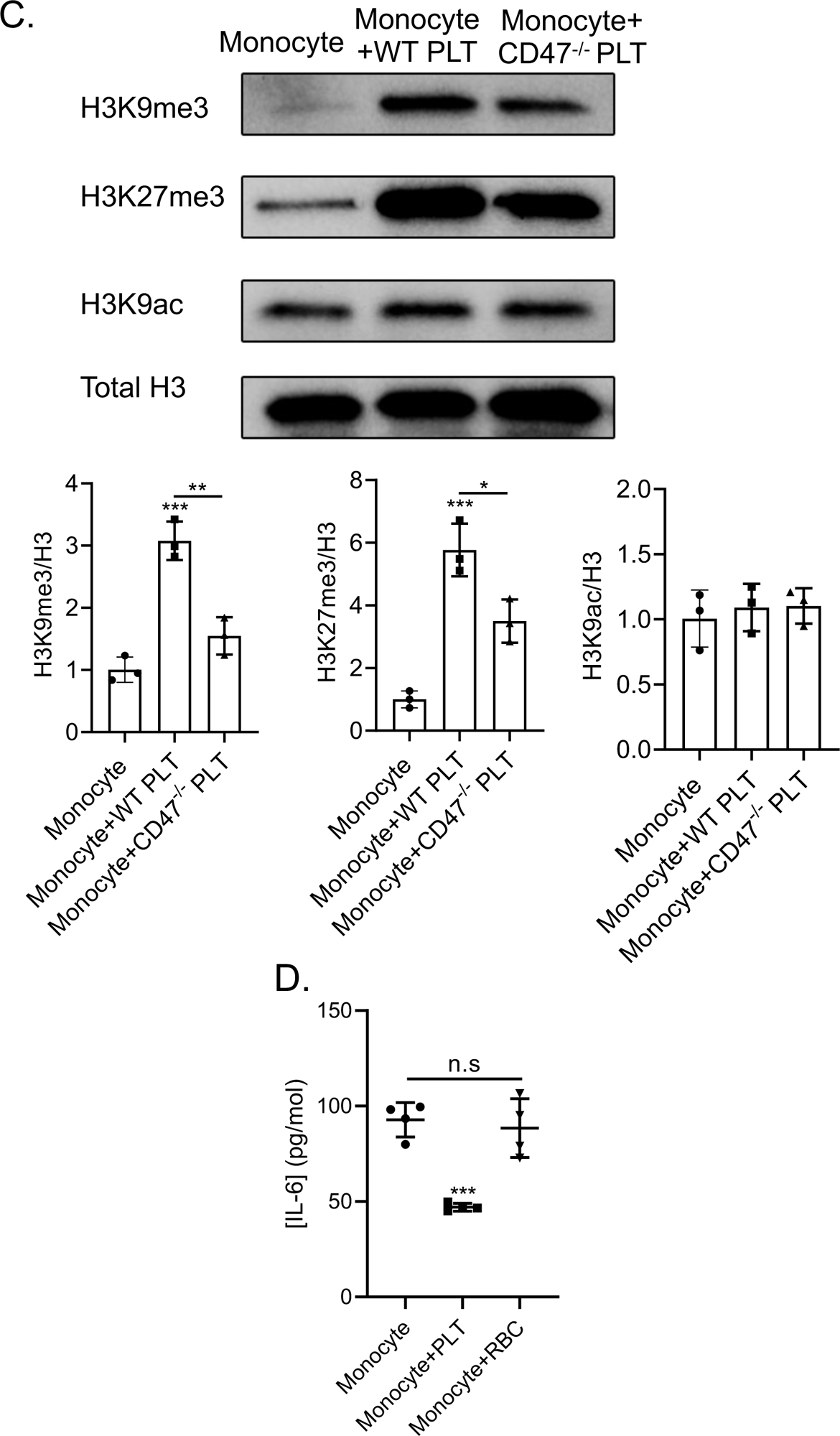
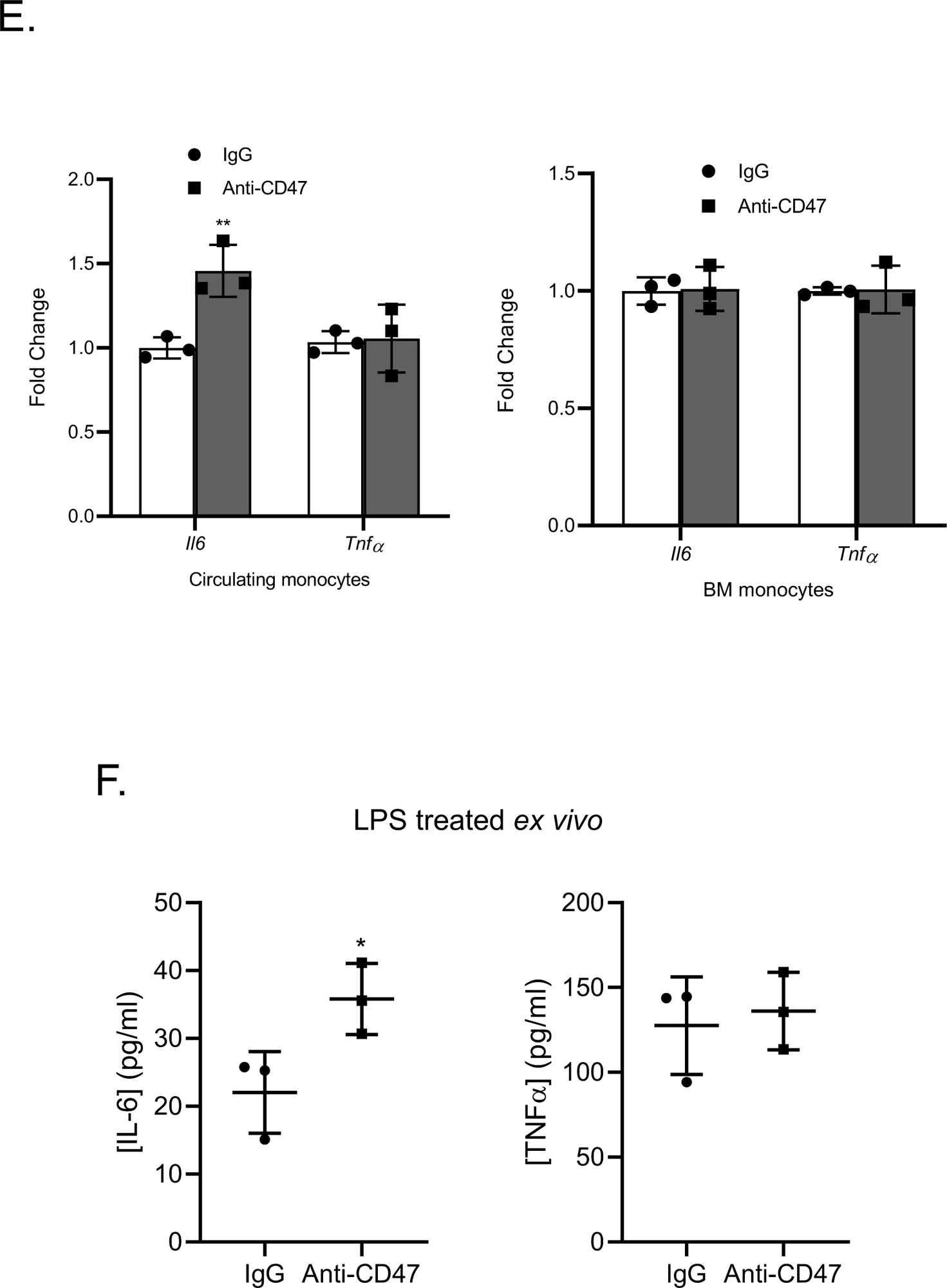
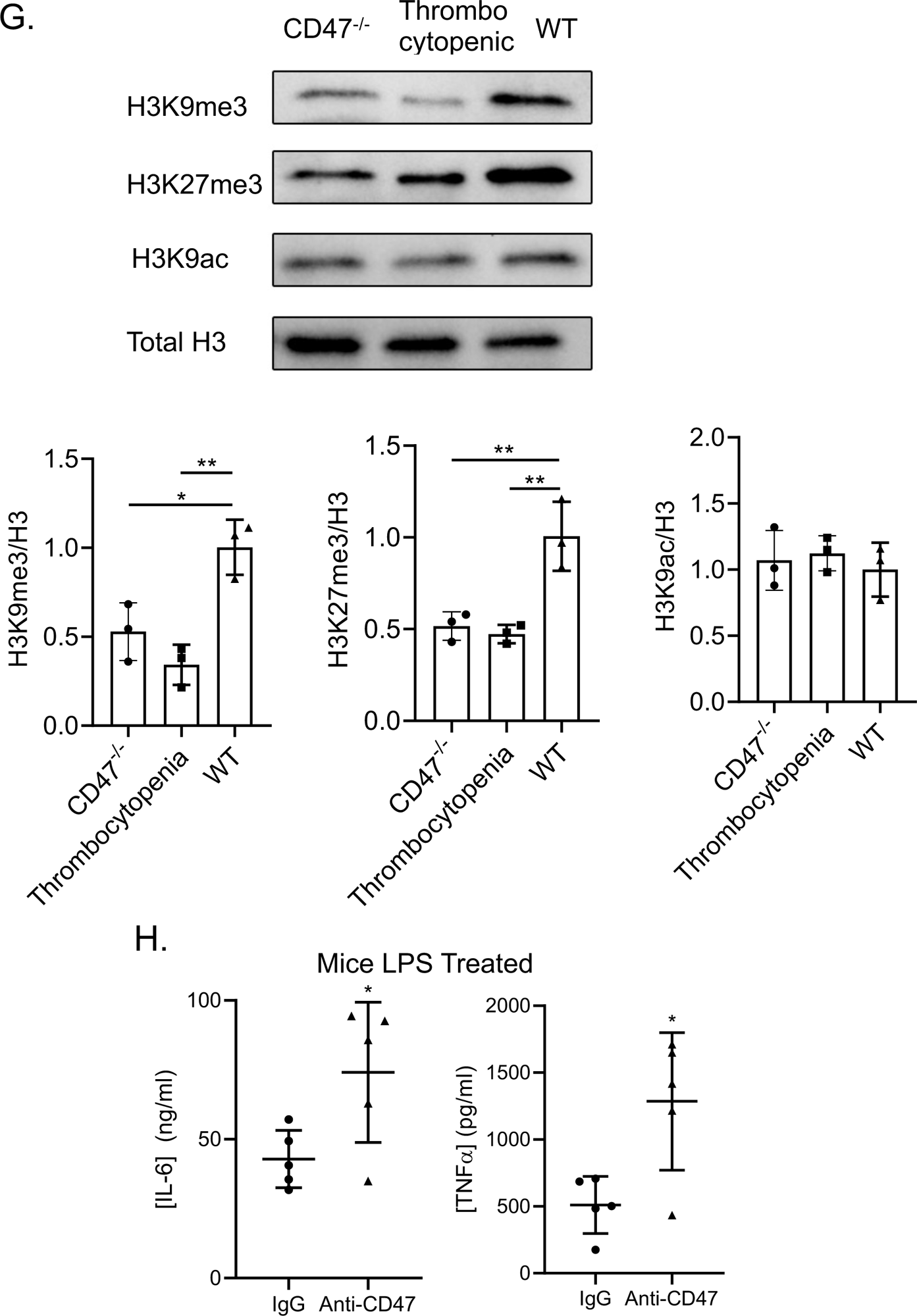
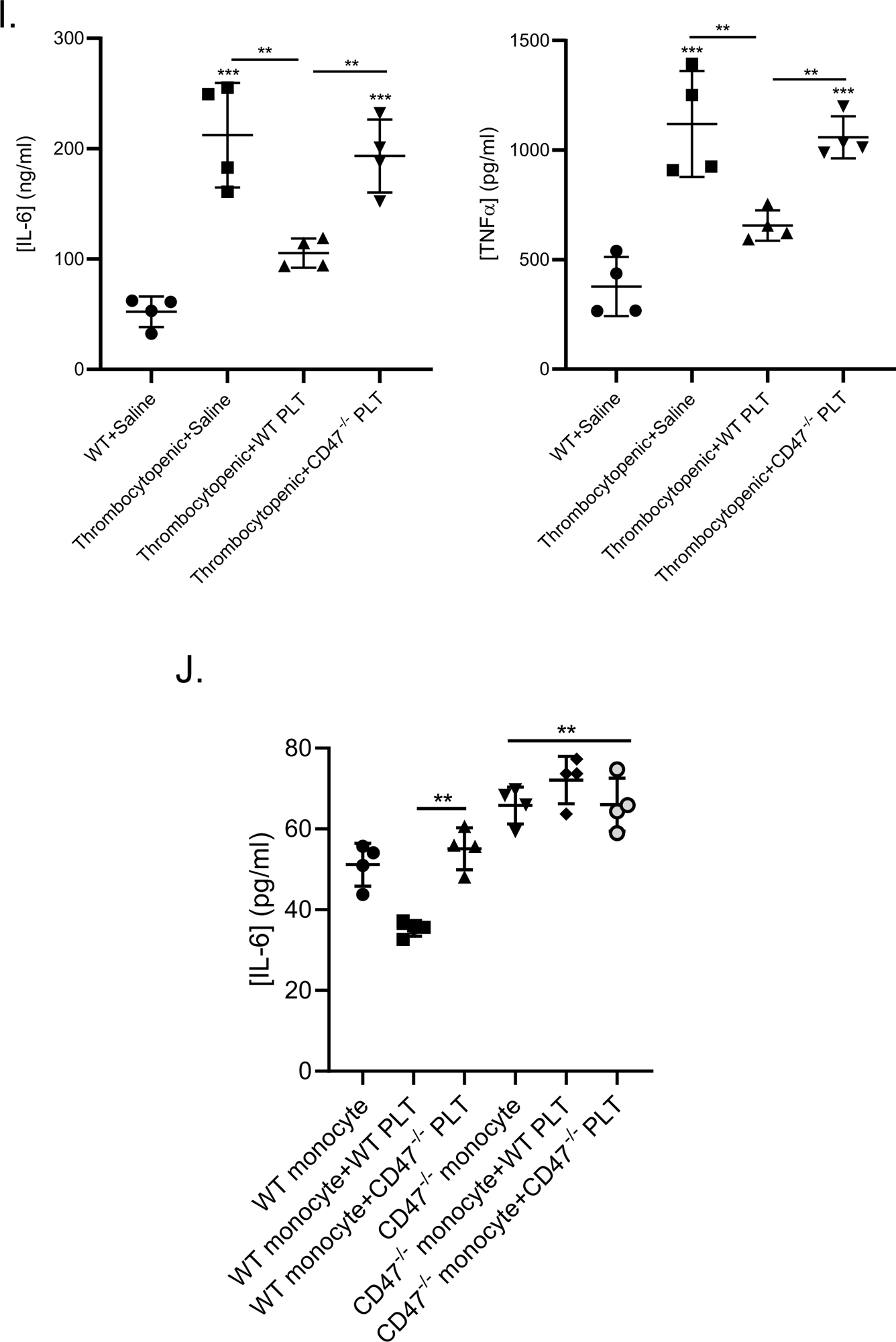
CD47 mediates platelet induced monocyte immune tolerance. A) Platelet mediated LPS tolerance is contact dependent. Monocytes or BMDM were cultured with platelets together or separately in a transwell chamber overnight and monocytes then stimulated with LPS. (*P < 0.05, **P < 0.01, ***P < 0.0001, by 1-way ANOVA followed by Tukey’s post hoc test). B) Platelet mediated monocyte immune tolerance is platelet CD47 dependent *in vitro.* Monocytes were incubated with RGDS treated or anti-CD47 Ab treated platelets, CD47^-/-^ platelets overnight and then LPS stimulated after platelets washed away. (*P < 0.05, **P < 0.01, by 1-way ANOVA followed by Tukey’s post hoc test). C) H3K9me3, H3K27me3 and H3K9ac immunoblots of monocytes cultured with WT or CD47^-/-^ platelets. (*P<0.05, **P<0.01, by 1-way ANOVA followed by Tukey’s post hoc test). D) Monocytes were incubated with platelets or RBCs overnight and then LPS stimulated. E-F) Platelet mediated monocyte immune tolerance is CD47 dependent *in vivo*. Mice were treated with anti-CD47 Ab or control IgG on d0, d2, and d4. On d5, peripheral or BM monocytes were isolated for E) qRT-PCR, and F) circulating monocytes were stimulated with LPS *ex vivo*. (*P < 0.05, **P < 0.01, by unpaired, 2-tailed Student’s *t* test). G) H3K9me3, H3K27me3 and H3K9ac immunoblots of circulating monocytes from CD47^-/-^, thrombocytopenic and WT mice. (*P<0.05, **P<0.01, by 1-way ANOVA followed by Tukey’s post hoc test). H) Control IgG and CD47 blocking Ab treated mice were LPS stimulated *in vivo*. (*P<0.05 by unpaired, 2-tailed Student’s *t* test). I) Transfusion of WT, but not CD47^-/-^ platelets to thrombocytopenic mice limited LPS induced cytokine production *in vivo*. (**P<0.01, ***P<0.0001, by 1-way ANOVA followed by Tukey’s post hoc test). J) Platelet CD47 mediated immune tolerance is dependent on platelet CD47 and monocyte CD47 interactions. WT platelet induced WT monocyte but not CD47^-/-^ monocyte LPS tolerance. (*P < 0.05, **P<0.01, ***P<0.0001, by 1-way ANOVA followed by Tukey’s post hoc test). Data shown as mean ± SEM.

Platelets interact with immune cells in both contact dependent and independent (secreted molecule) manners (18, 49, 50). To determine whether platelets limit monocyte LPS responses in a cell-contact dependent or independent manner, transwell chambers were used to separate platelets from BM monocytes or BMDM. When platelets were physically separated from monocytes or BMDM, platelets no longer limited LPS induced IL-6 production (**Fig 4A**). Furthermore, incubation of monocytes with both resting platelets or activated and washed platelets, limited monocyte IL-6 secretion, but this response was not noted following incubation with activated platelet releasates or platelet conditioned media (**Supplemental Fig 4A**). These results highlight that direct platelet-monocyte contact is needed for platelet mediated monocyte LPS tolerance, while activated platelet releasates do not, and further demonstrate the contact specificity in our *in vitro* model system.

Resting platelets express surface proteins such as GPIbα and GPVI, and activated platelets increase surface proteins such as P-selectin and CD40L, that mediate direct interactions with leukocytes to regulate immune cell migration, differentiation and cytokine production (50). Platelet limited monocyte IL-6 in response to LPS was not altered when co-cultures were performed in the presence of P-selectin or CD40L blocking antibodies, nor in the presence of the integrin blocker RGDS (**Fig 4B** **and Supplemental Fig 4B**). PSGL-1 and Mac-1 complex (CD11b and CD18) are counter-receptors on monocytes for P-selectin and GPIb respectively. Monocytes treated with PSGL-1, CD11b or CD18 blocking antibody in co-culture produced similar amounts of IL-6 in response to LPS compared to IgG treated monocytes (**Supplemental Fig 4C**). We also verified that platelet induced monocyte responses were not dependent on platelet phosphatidylserine, thrombopoietin (TPO) or TGFβ (**Supplemental Fig 4C**). Resting platelets also express CD47, and when platelets and monocytes were cultured with CD47 blocking antibody, platelets no longer attenuated LPS induced IL-6 (**Fig 4B**). Similarly, monocytes incubated with CD47^-/-^ platelets did not have an attenuated LPS responses (**Fig 4B**). Monocytes formed similar aggregates with CD47^-/-^ platelets after overnight co-culture compared to WT platelets (**Supplemental Fig 4D**). However, histone H3K9me3 and H3K27me3 levels were diminished when monocytes were co-cultured with CD47^-/-^ platelets (**Fig 4C**).

CD47 is an immunoglobulin (Ig) superfamily membrane protein expressed on hematopoietic and non-hematopoietic cells. It serves as a receptor for thrombospondin (TSP) and a ligand for the SIRPα receptor to regulate cell adhesion/migration, apoptosis and phagocytosis (51–53). CD47 is most recognized for delivering ‘don’t eat me’ signals and regulating red blood cell (RBC) clearance. CD47 is overexpressed on surface of many cancer cells to inhibit macrophage mediated phagocytosis and anti-CD47 monoclonal antibodies are currently in clinical trials for both leukemia and solid tumors (54). However, CD47 has cell signaling roles and CD47^-/-^ mice have minimal changes in platelet count (55) (**Supplemental Figure 5**). Unlike platelets, RBC incubation with monocytes did not limit LPS induced IL-6 (**Fig 4D**), indicating platelet CD47 has unique cell tolerance signaling roles that are different from RBC expressed CD47. To investigate the role of CD47 in LPS tolerance *in vivo*, we treated mice with CD47 blocking antibody or control IgG. Similar to thrombocytopenic mice, circulating monocytes isolated from CD47 blocking antibody treated mice had increased basal *Il6* and secreted more IL-6 in response to LPS *ex vivo*, compared to IgG treated mice (**Fig 4E-F**). These data were recapitulated in circulating monocytes isolated from CD47^-/-^ mice (**Supplemental Fig 4E**). Immunoblots showed circulating monocytes from CD47^-/-^ mice had similar levels of H3K9me3 and H3K27me3 compared to monocytes from thrombocytopenic mice, while monocytes from WT mice had greater histone methylation (**Fig 4G**). Mice treated with CD47 blocking antibody prior to LPS challenge also had significantly higher plasma IL-6 and TNFα *in vivo* (**Fig 4H**). These data demonstrate that platelet CD47 delivers an LPS tolerant signal to monocytes through histone methylation both *in vitro* and *in vivo*. To demonstrate platelet CD47 specificity *in vivo* we performed platelet transfusion experiments in which thrombocytopenic mice were transfused with either saline, washed WT platelets, or CD47^-/-^ platelets on d4 post-DT, and 24 hrs later mice were given LPS. Plasma IL-6 and TNFα concentrations were higher in thrombocytopenic mice compared to control mice, and this was largely reversed by the transfusion of WT platelets (**Fig 4I**). However, CD47^-/-^ platelet transfusions did not significantly alter plasma IL-6 or TNFα compared to saline treated thrombocytopenic mice (**Fig 4I**). These results suggest that platelet mediated monocyte immune tolerance is dependent on platelet CD47.

Most studies of CD47 interactions with monocytes have focused on SIRPα that delivers an anti-phagocytic signal to monocytes. To determine whether monocyte SIRPα is the platelet CD47 receptor that contributes to immune regulation, we treated monocytes with SIRPα blocking antibody or a SHP (downstream effector of SIRPα) inhibitor (PTP I inhibitor) prior to platelet-monocyte co-culture. Neither SIRPα blocking, nor SHP inhibition, reversed platelet mediated LPS signaling inhibition (**Supplemental Fig 4F**). CD47 can form homotypic interactions with CD47 on other cells to elicit CD47 signal transduction(56, 57). For this reason, and given that monocytes also express CD47, we evaluated CD47-CD47 signaling using CD47^-/-^ platelets and CD47^-/-^ monocytes. As prior, WT platelets, not CD47^-/-^ platelets suppressed WT monocyte LPS induced IL-6 production. However, WT platelet mediated monocyte inhibition was abrogated when co-cultured with CD47^-/-^ monocytes, indicating the essential role of monocyte CD47 in platelet mediated LPS tolerance (**Fig 4J**). Together, these data suggest that platelet-monocyte CD47 homotypic interactions are crucial to platelet mediated monocyte immune tolerance.

To gain a more complete concept of resting platelet regulating monocyte responses both before and after LPS stimulation, we performed RNA-Seq on mRNA isolated from monocytes incubated in control media (NC) or with platelets overnight (PLT), as well as mRNA isolated from control monocytes 4 hrs after stimulation with LPS (LPS) or mRNA isolated from LPS stimulated monocytes co-incubated with platelets (PLT LPS). Platelet only treated monocytes exhibited different transcription patterns compared to control media treated monocytes, both before and after LPS treatment (**Fig 5A**). Gene ontology (GO) enrichment analysis of genes that were upregulated by platelet interactions revealed significantly enriched sets of glucose metabolism/glycolysis and immune regulatory pathways (**Fig 5B**), while gene pathways suppressed by platelets included immune response pathways such as responses to bacteria origin/LPS, consistent with resting platelet induced monocyte LPS tolerant responses (**Fig 5B**). Similarly, 4 hours after LPS stimulation, prior platelet co-culture tended to amplify monocyte gene expression involved in cell cycle/proliferation and glycolytic processes, while decreasing the expression of genes related to innate immune signaling pathways (**Fig 5B**). Among the 32 genes related to LPS response/TLR related pathway identified in RNA-Seq, 17, including *Il6* and *Il1b*, had chromatin regions that were significantly more accessible in monocytes from thrombocytopenic mice (**Fig 5C**). This suggests that chromatin remodeling is at least in part responsible for the transcriptional signature of LPS response pathways. RNA-Seq data was further confirmed by qRT-PCR and platelet co-cultured monocytes had higher expression of glycolytic genes including *Ldha*, *Pfkl*, *Tpi1*, *Hif1α* and lower expression of inflammatory genes including *Il6* and *Il1b* compared to control monocytes (**Fig 5D**). Consistent with the RNA-seq data, prior exposure of monocytes to platelets primed the production of IL-10 and CCL2, but limited CXCL1 (**Fig 5E-F**), further demonstrating resting platelet mediated monocyte reprogramming, rather than a complete response suppression. RNA-seq pathways regulated by resting platelets included bacteria responses. Prior platelet co-culture attenuated monocyte Pam2CSK4 (TLR2 agonist) and CpG (TLR9 agonist) induced responses, but had little effect on IL1β induced responses, indicating platelet mediated monocyte immune tolerant response regulation is more TLR skewed (**Fig 5F**). Taken together, these results indicated that incubation of monocytes with platelets led to an enhanced glycolysis gene expression and TLR tolerant immune responses.

**Figure 5.**
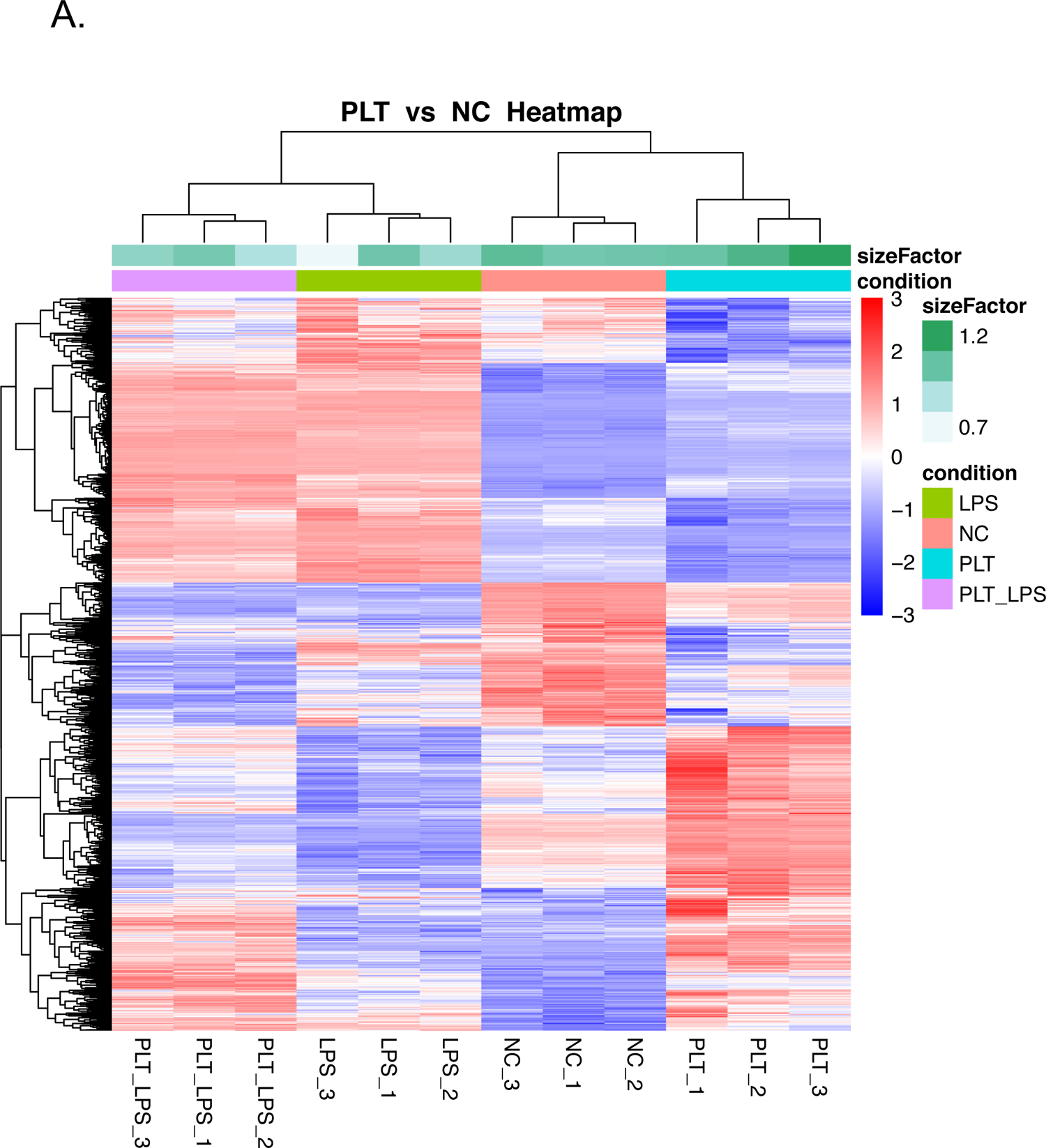
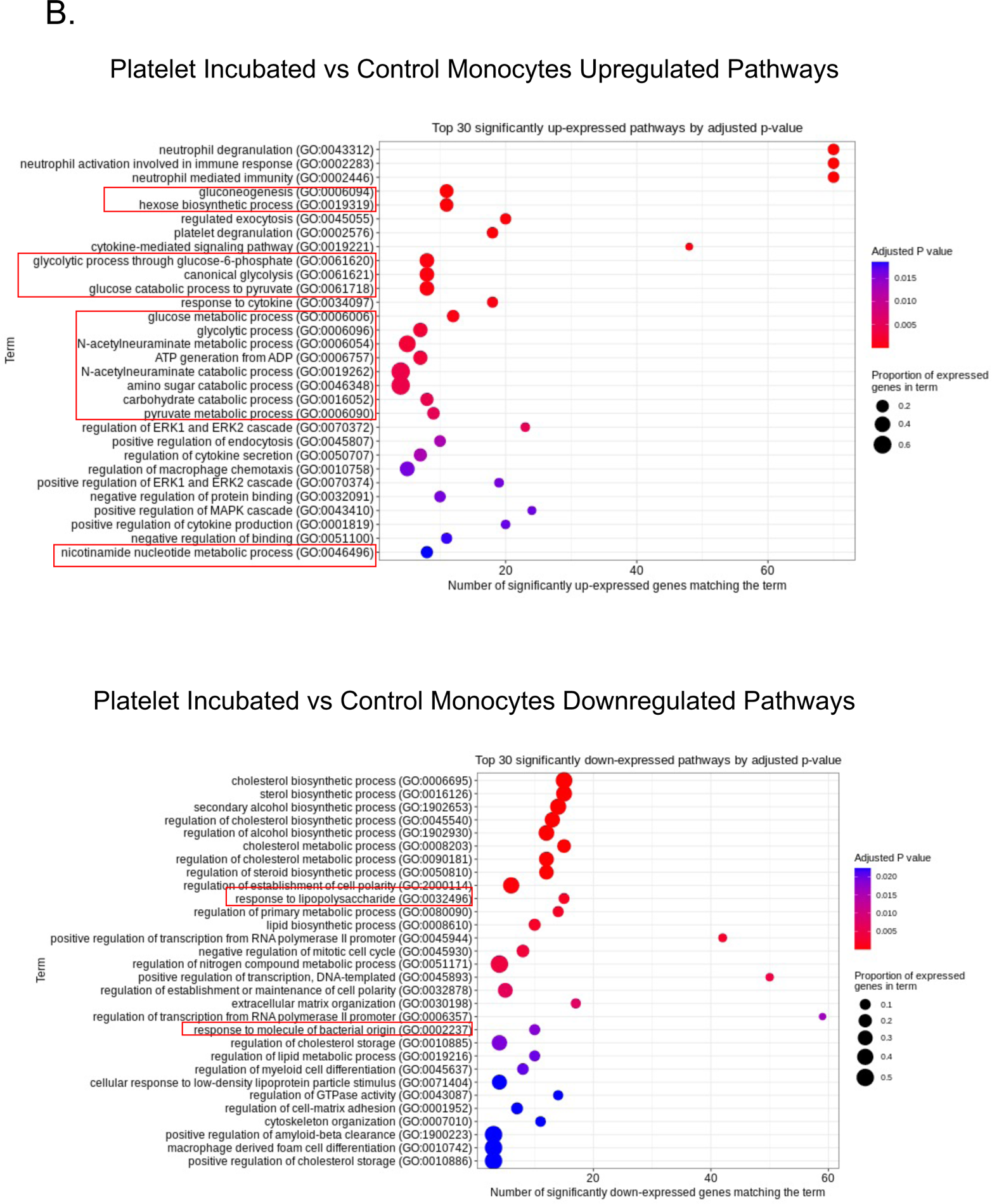
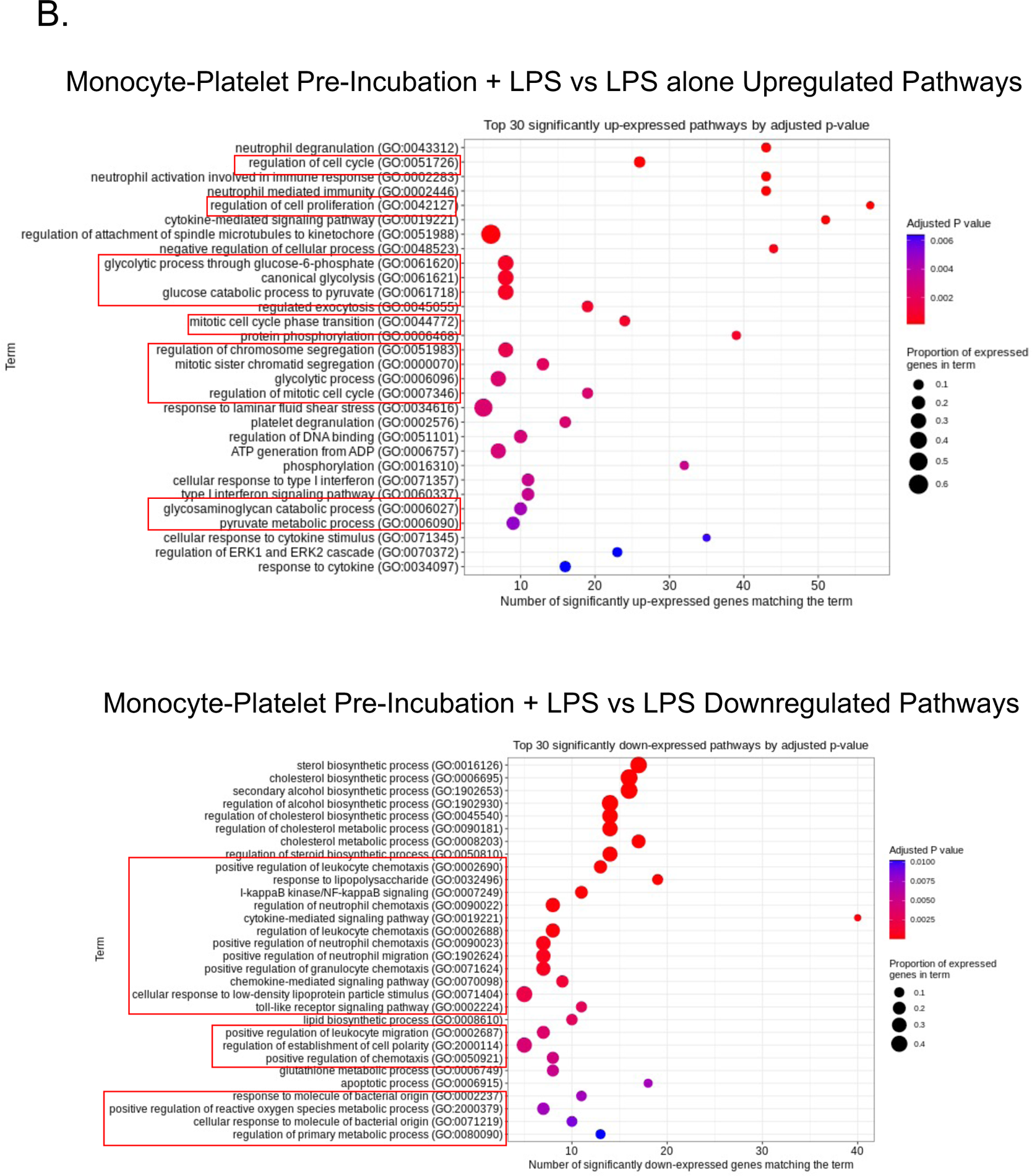
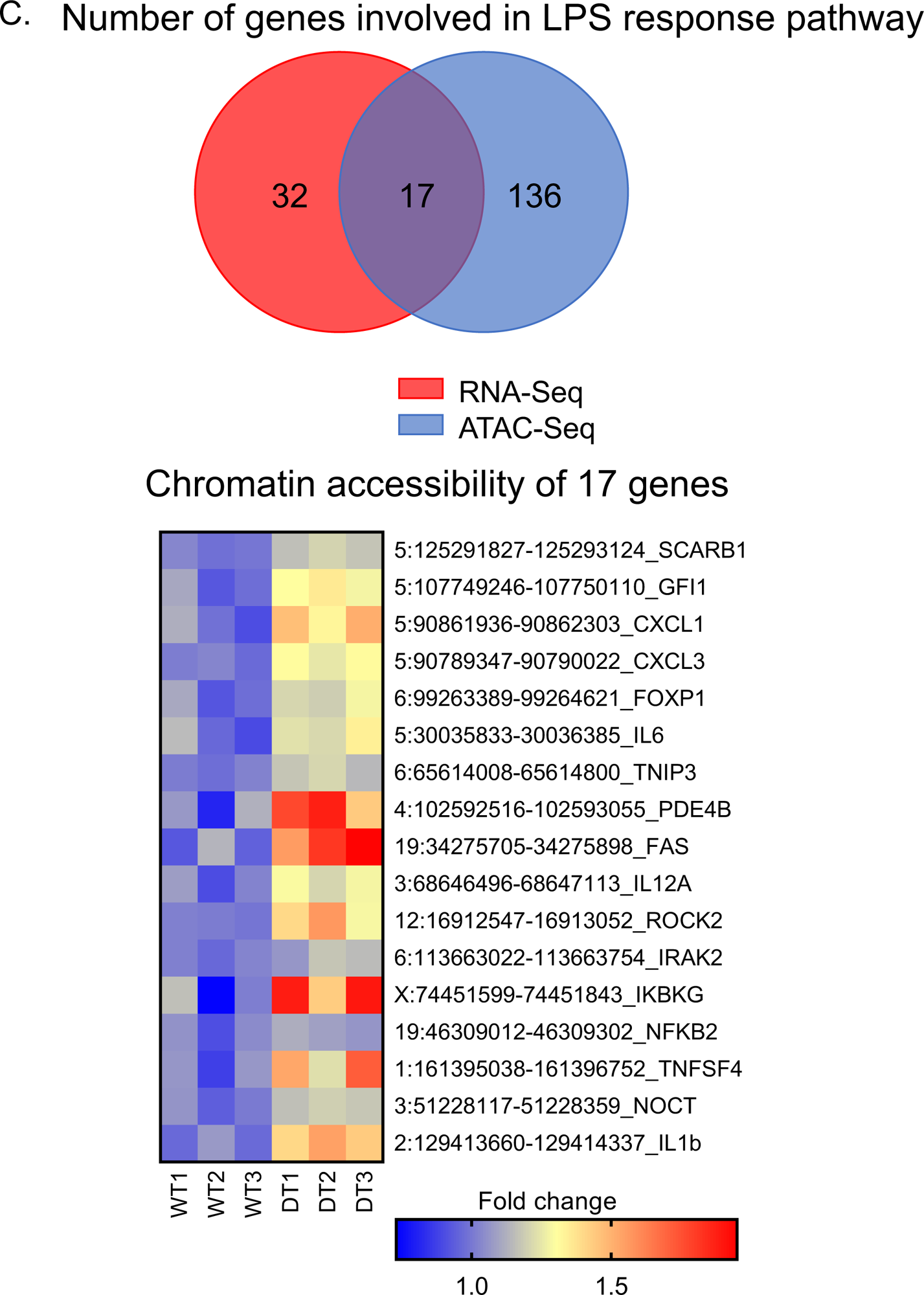
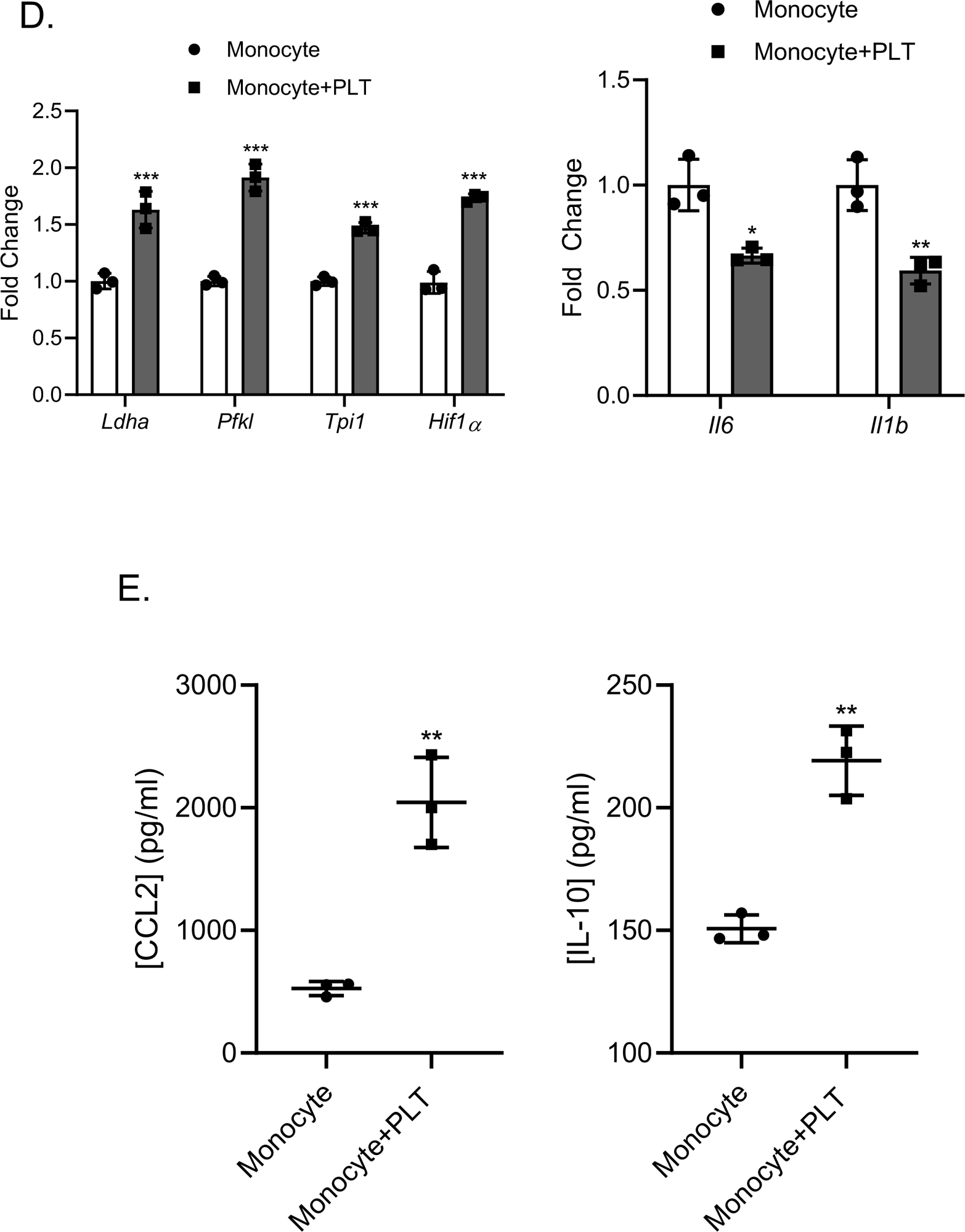
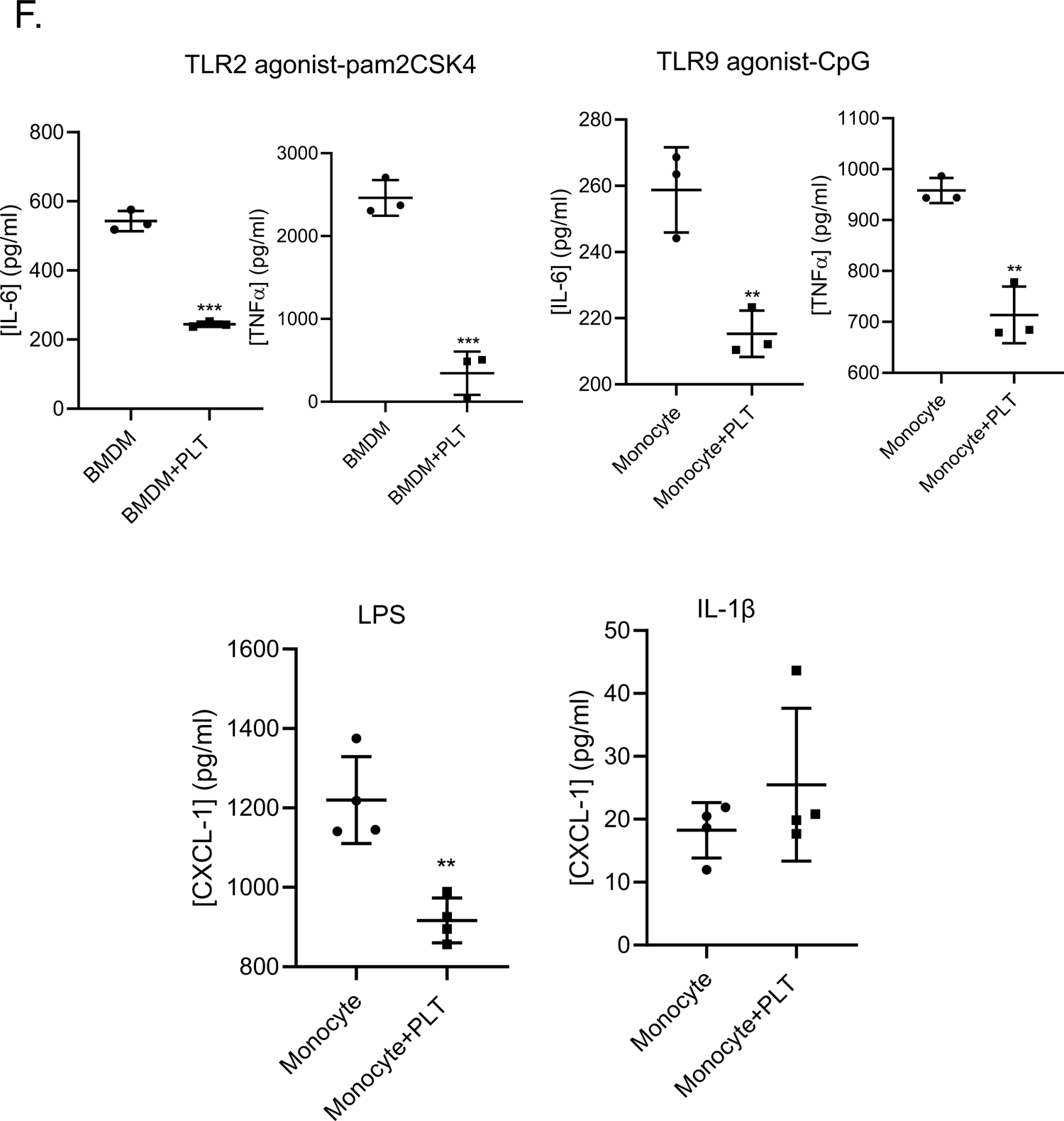
Platelets induce monocyte metabolic pathways and limit TLR signaling. A-B) Monocytes were incubated overnight in control buffer or with platelets. Cells were washed to remove platelets and mRNA isolated for RNA-seq or monocytes were LPS stimulated and 4 hrs later mRNA was isolated for RNA-seq. A) Platelet incubated monocytes had different gene expression patterns. B) GO analysis of genes regulated by platelets. Platelets regulated monocyte metabolism and immune related gene pathways. C) Venn diagrams showing the intersection between changed LPS response genes by platelets in RNA-Seq and ATAC-Seq. Heatmap showing normalized accessibility fold change of those 17 LPS response associated gene regions. D) Confirmation of metabolism related genes and immune related genes by D) qRT-PCR and E) ELISA. Monocytes were incubated with control buffer or platelets overnight and qRT-PCR/ELISA performed. (*P<0.05, **P<0.01, ***P<0.001, by unpaired, 2-tailed Student’s *t* test). F) Monocyte/macrophage responses to multiple TLR associated ligands is reduced by platelet pre-incubation, but IL-1β is not. (**P < 0.01, ***P < 0.001, by unpaired, 2-tailed Student’s *t* test). Data shown as mean ± SEM

To functionally validate monocyte glycolysis induction by platelets, Seahorse glycolytic rate assays were performed on circulating monocytes isolated from WT control and thrombocytopenic mice. Monocytes isolated from TPOR^-/-^ mice had lower extracellular acidification rate (ECAR) in both basal conditions (basal glycolysis) and after rotenone/antimycin A treatment (compensatory glycolysis) compared to WT mouse isolated monocytes (**Fig 6A**). Similar results were found in acutely thrombocytopenic mice, further suggesting platelets regulate monocyte glycolysis *in vivo* (**Fig 6A**). Co-culture of monocytes with WT platelets *in vitro* increased monocyte glycolytic rate, but this effect was limited using CD47 blocking antibody treated platelets or platelets from CD47^-/-^ mice (**Supplemental Fig 6**, note some platelets adhere to the Seahorse plate after washing resulting in partial reversal with CD47^-/-^ platelets). Together, this highlights the contribution of platelets in enhancing monocyte glycolysis. Metabolic reprogramming of immune cells regulates cell phenotype, functional plasticity, and epigenetic modifications that are critical mediators of innate immune training (58–60). We therefore hypothesized that platelet programmed monocyte metabolism toward glycolysis is essential to platelet mediated immune tolerance. To explore this, we inhibited monocyte glycolysis using either glucose-free media or 2-deoxy-glucose (2-DG) in platelet-monocyte co-cultures. Monocyte tolerant responses induced by platelets was abrogated when glycolysis was inhibited, demonstrating the crucial role of platelet mediated glycolysis in mounting LPS immune tolerance (**Fig 6B**). Similarly, in glucose free media, platelets no longer induced histone methylation (**Fig 6C**), further indicating that platelets mediate histone methylation in a glycolysis dependent manner.

**Figure 6.**
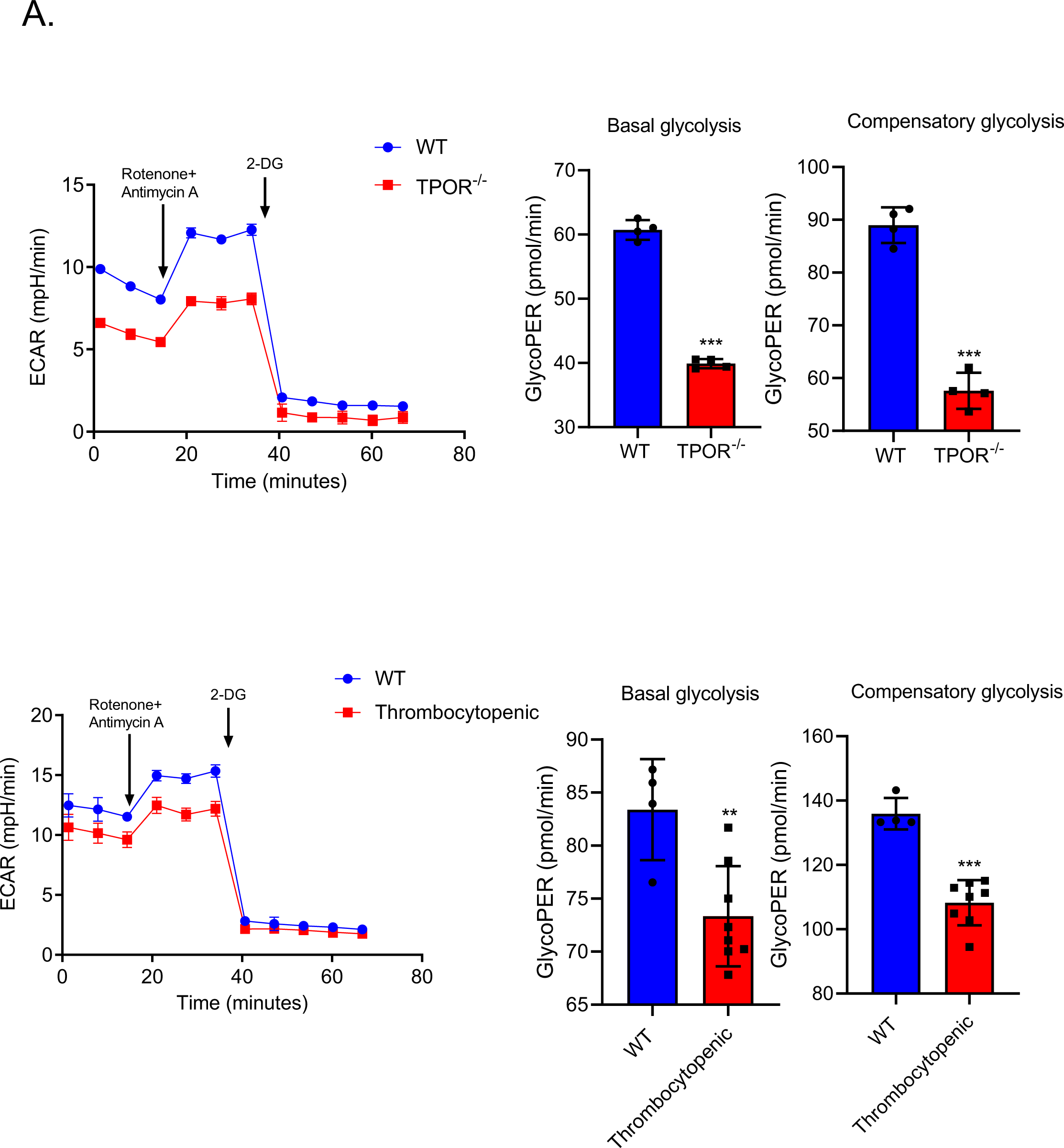
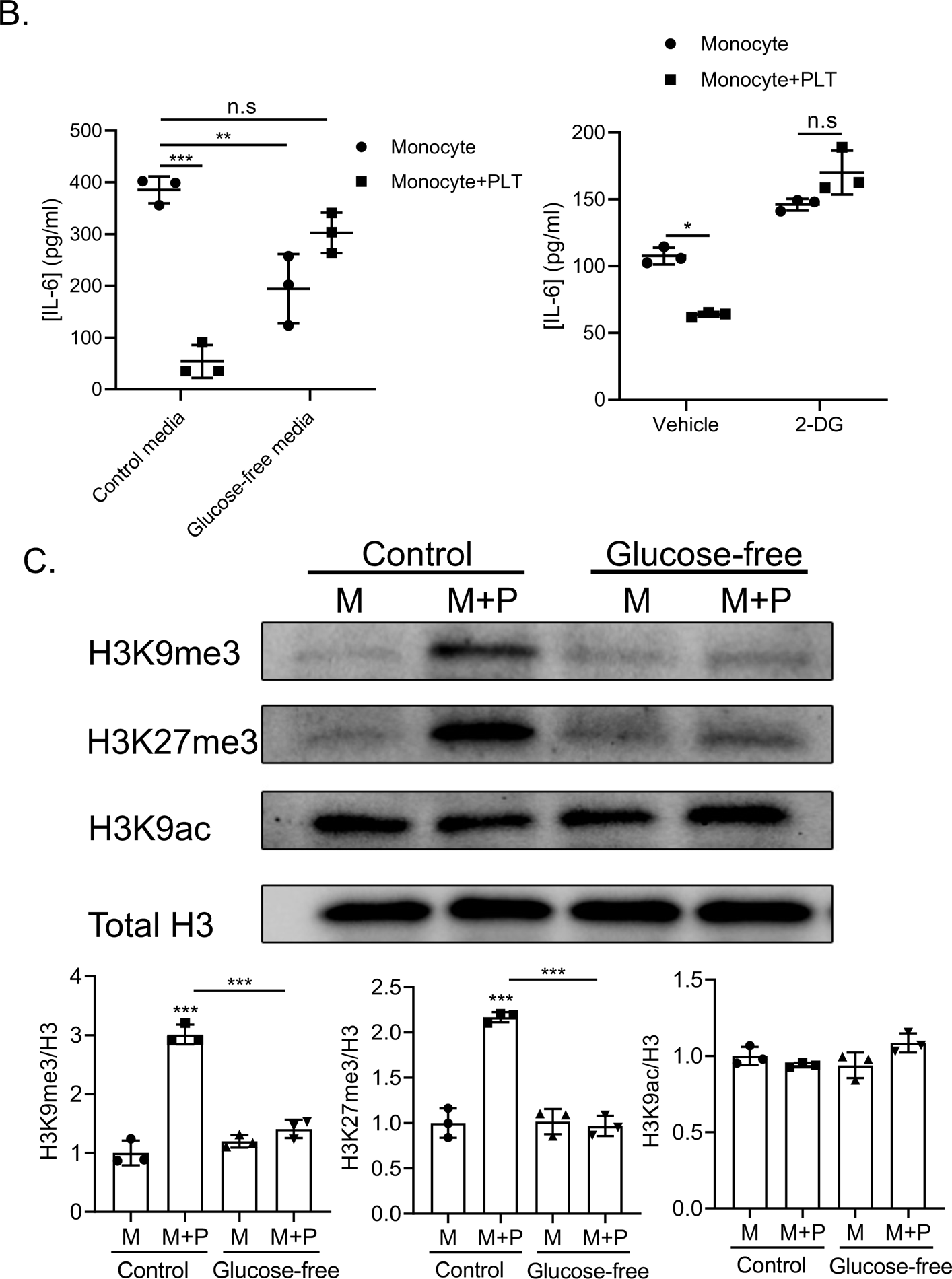
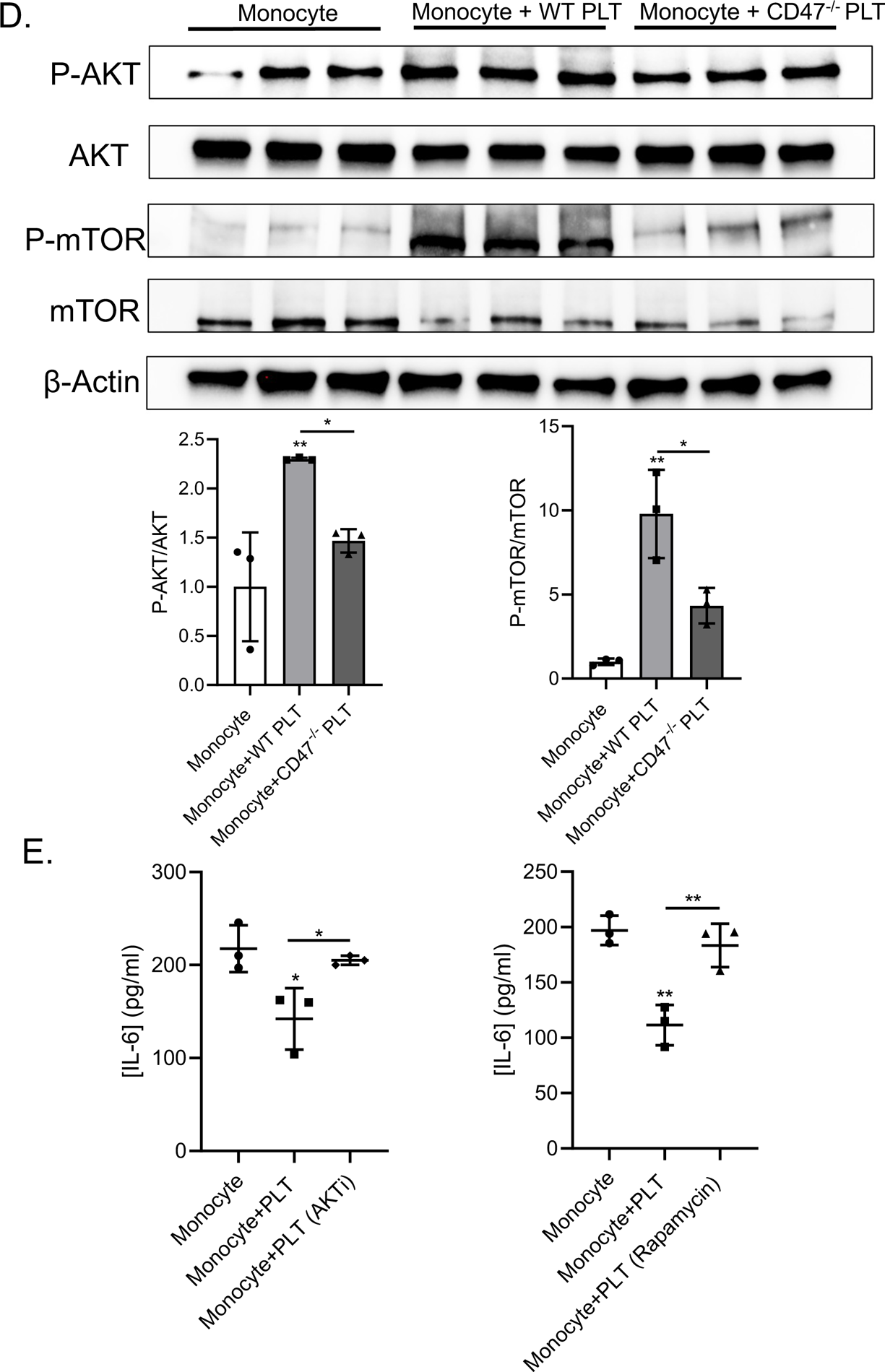
Platelets induce monocyte glycolysis as a mechanism of trained tolerance. A) Monocytes from thrombocytopenic mice had reduced basal and compensatory glycolysis. Glycolytic rate assay was performed on monocytes isolated from WT, TPOR^-/-^ mice, or DT induced thrombocytopenic mice on d5 post-DT. (**P<0.01, ***P<0.001, by unpaired, 2-tailed Student’s *t* test). B) Platelets and monocytes co-cultured in glucose free or 2-DG media had no change in IL-6 in response to LPS. Monocytes were treated with 2-DG prior to platelet co-culture or co-culture was performed in glucose free media and then cells LPS stimulated. (*P < 0.05, **P<0.01, ***P<0.0001, by 2-way ANOVA followed by Tukey’s post hoc test). C) H3K9me3, H3K27me3 and H3K9ac immunoblots of monocytes incubated with platelets in control or glucose free media (M=monocyte, M+P=monocyte/platelet co-cultures. ***P<0.001, by 1-way ANOVA followed by Tukey’s post hoc test). D) Platelets induced monocyte AKT and mTOR phosphorylation. P-mTOR and P-AKT immunoblots of monocytes co-cultured with WT or CD47^-/-^ platelets. (*P<0.05, **P<0.01, by 1-way ANOVA followed by Tukey’s post hoc test). E) Platelet mediated monocyte immune training is AKT/mTOR dependent. Monocytes were treated with mTOR or AKT inhibitor prior to platelet co-culture and then cells LPS stimulated. (*P<0.05, **P<0.01, by 1-way ANOVA followed by Tukey’s post hoc test). Data shown as mean ± SEM.

mTOR acts as a metabolomic environment sensor and functions in innate immune cell development, polarization, and cytokine production, and AKT/mTOR signaling is a master regulator of glucose metabolism and innate immune training (61–63). Monocytes incubated with WT platelets demonstrated significant activation of both mTOR and AKT (**Fig 6D**). Both mTOR and AKT activation are at least partially dependent on platelet CD47, demonstrated by monocytes incubated with CD47^-/-^ platelets having significantly less mTOR and AKT phosphorylation compared to WT platelet co-incubated monocytes (**Fig 6D**). To determine the causality between AKT/mTOR pathway activation and platelet mediated immune training, monocyte mTOR was inhibited using rapamycin during platelet co-culture which limited platelet induced LPS tolerance (**Fig 6E**). The AKT inhibitor MK2206 similarly blocked platelet induced monocyte immune tolerance (**Fig 6E**), demonstrating the relationship between platelet CD47 dependent activation of the AKT/mTOR pathway and immune tolerance in platelet trained monocytes.

Together, these data demonstrate that circulating resting platelets limit monocyte LPS responses in normal conditions, in a manner that is dependent on CD47 mediated, metabolism driven, epigenetic programming (**Fig 7**).

**Figure 7.**
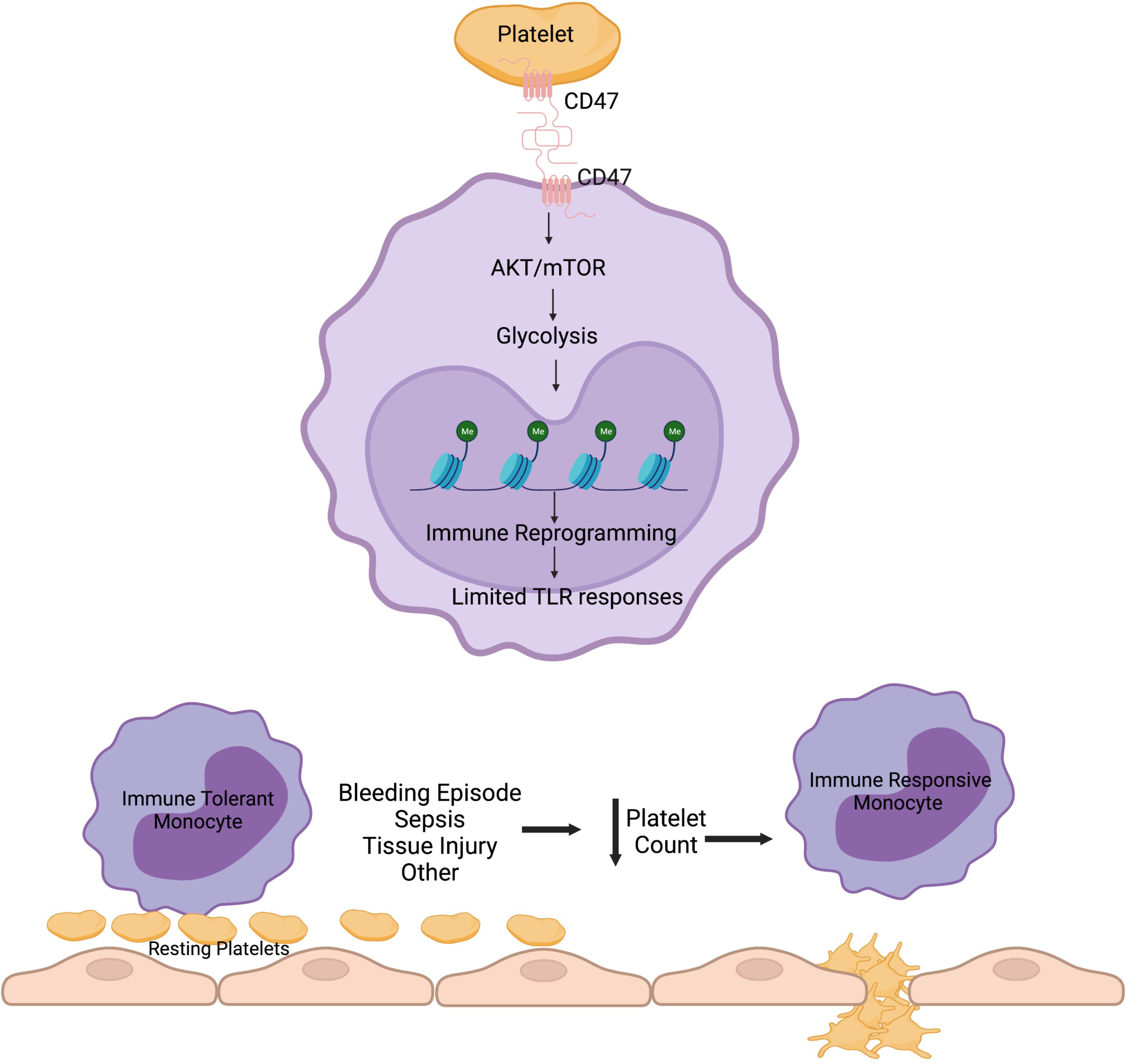
Data Summary. Platelets regulate monocyte immune programming in a manner that is dependent on CD47-CD47 interactions that induces monocyte metabolism, epigenetic remodeling and immune programming. Therefore, a decline in platelet count leads to monocyte immune dysfunction.

## Discussion

We have discovered that resting platelet interactions with circulating monocytes, induces monocyte glycolysis, which is in turn crucial for chromatin remodeling and the maintenance of monocyte immune tolerance. Our studies indicate that circulating platelets in normal healthy conditions limit monocyte immune dysregulation, and therefore a decline in platelet count alters monocyte responses, independent of the cause of thrombocytopenia (**Fig 7**). This makes physiologic sense: thrombocytopenia is indicative of significant vascular injury or infection and signals the need for heightened immune responses, making platelet numbers a valuable immune biosensor. Work from our research group has found that platelets not only have roles in promoting immune responses at times of tissue injury or infection, but platelets also have central roles in maintaining steady-state immune homeostasis. For example, chronically thrombocytopenic mice, even in the absence of any immune stimulation, had increased numbers of Th17 type of T helper cells, demonstrating a need for platelets to limit dysfunctional T cell differentiation (64). Platelets interact with immune cells in both contact dependent and independent manners, and platelets circulate at the interface between the vessel wall and immune cells, positioning platelets as sentinels of vascular injury and inflammation (65, 66). Results of this study lead us to propose the novel concept, that in healthy conditions platelets limit immune cell activation and promote a more quiescent immune environment. This means that activated platelets induce immune cell activation, while a decline in platelet count provides a further physiologic cue of the need to heighten immune responses. Thrombocytopenia therefore leads to a propensity for immune dysfunction, independent of the cause of the platelet decline.

These data provide novel insights into prior unknown roles for platelets in immune homeostasis, and a novel discovery of endogenous cell based innate immune training in basal conditions. In the context of sepsis, this means that infection induced platelet activation early post-infection may drive monocyte activation that is then exacerbated by a rapid decline in platelet count and further monocyte dysfunction. Monocytes are consumed or tissue migrate early in infection but continue to be rapidly produced from the bone marrow, perhaps amplifying the inflammatory priming of monocytes as they emerge more rapidly from the marrow into thrombocytopenic conditions (67). Therefore, strategies to limit sepsis associated thrombocytopenia may help to limit the associated immune dysfunction. Thrombocytopenia is also seen in about 20% of acute coronary syndrome (ACS) patients, up to 50% of patients put on extracorporeal membrane oxygenation (EMCO), as well as in other infectious diseases such as malaria, and is similarly associated with increased risk of inflammatory complications and mortality (68–73). The results of our studies may therefore be more broadly generalizable and lead to a deeper mechanistic understanding of immune dysfunction in many disease states.

Innate immune training continues to be conceptually defined. This is the first demonstration that platelets regulate monocyte metabolism and gene programming in healthy conditions and places a sterile cell-cell interaction function for platelets in immune homeostasis, in addition to prior studied roles for activated platelets in driving immune responses. It is interesting to note that platelet releasates and intact platelets in co-culture with monocytes induced different monocyte responses to LPS. This leads to a concept that activated platelet derived products promote acute monocyte inflammatory responses, while in the absence of platelet activation, platelet-monocyte interactions lead to monocyte immune tolerance. There are many well described mediators of activated platelet induced monocyte inflammation, and activated platelets secrete many products that increase monocyte inflammatory responses, including our past studies demonstrating mechanisms of PF4 and b2M induced monocyte inflammatory activation (74, 75). Whether resting circulating platelets also mediate immune quiescence in other cells is not known, but thrombocytopenia does lead to endothelial leak, but platelet dependent endothelial cell reprogramming is yet to be defined.

CD47 is best known for delivering ‘don’t eat me signals’ that limit phagocytosis, and a decline in CD47 as RBCs age provides a physiologic signal to clear aged RBCs, but platelet CD47 does not have a major role in regulating platelet lifespans (55). CD47 has less studied cell signaling functions that is in part dependent on its receptor complex (76–78). CD47 was also referred to as integrin associated protein (IAP) because it is found in a complex with integrins primarily of the β1 or β3 family (*α*_2_*β*_1_ and *α*II_b_*β* _3_ integrins), but integrin blocking did not affect platelet induced LPS tolerance (**Fig 4B**). In addition to delivering don’t eat me signals, CD47 may regulate cell functions such as apoptosis, AKT dependent proliferation, cell adhesion, and cell migration. Incubating monocytes with RBCs did not alter LPS responses, indicating that platelet CD47 has a unique monocyte regulation function that RBC CD47 does not. CD47 complexes can vary significantly between cell types and CD47 forms a cell membrane complex with FasR on T Cells, CD14 and TLR4 on innate immune cells, VEGFR on endothelial cells, and with CD36 and integrins on many cell types, as well as associating with lipid rafts and GPCR complexes(79, 80). CD47 can also be heavily glycosylated affecting its protein interactions and signaling potential and the cytoplasmic tail of CD47 is alternatively spliced in different cell types. These cell specific variables of CD47 expression and interactions may account for why platelet, but not RBC, CD47 modulates monocyte immune tolerance. Much further study is needed to define these unique platelet CD47 signaling functions.

## Method

### Mouse studies

All mice were on a C57BL6/J background. PF4-DTR mice were generated by crossing PF4-Cre mice (The Jackson Laboratory) with simian iDTR mice containing a LoxP-flanked stop sequence. Wild-type mice and CD47^-/-^ mice were obtained from Jackson Laboratory. TPOR^-/-^ mice were bred in house as described in our past studies (29). To obtain whole blood, mice were bled retro-orbital into EDTA blood collection tube (BD Microtainer tube).

Mice treated with anti-CD47 antibody (Clone MIAP410, Bio X Cell) received initial injection (200 μg) on day 0 and injections (200 μg) on day 2 and day 4 i.p. before LPS induced sepsis on day 5. Mouse peripheral blood was collected 4 hours after LPS challenge.

### LPS-induced sepsis

Mice (8-10 weeks old) were injected i.p. with 5 mg/kg LPS (*E coli* O55:B5, purchased from Sigma) in a volume of 100 μl of saline.

### Cecal slurry induced sepsis

Fresh feces were collected from the lower cecum of euthanized donor mice (8-12 weeks), weighed, and mixed with PBS containing 10% glycerol. Fecal slurry was vortexed and filtered through 70 μm strainer. Each mouse was given an i.p. injection of 0.8 mg/g fecal slurry. To ensure reproducibility, fresh fecal solution was prepared from mice living in the same conditions as the experimental animals and cecal slurry preps frozen at -80°C.

### Platelet depletion procedure

WT and PF4-DTR mice were injected i.p. with either 100 μl phosphate buffered saline (PBS) (Corning) or 400 ng diphtheria toxin (DT) (Cayman Chemical) on day 0. Blood was collected every day until day 9 in EDTA blood collection tube (BD Microtainer tube) to measure the platelet count using an Abaxis VetScan HM5. D3 and d5 blood were also collected for HMGB1 (Antibodies-online.com) and S100A8/A9 (DuoSet, R&D Systems) measurement by ELISA.

### Platelet studies

Mouse platelets were obtained by retro-orbital bleed into heparinized Tyrodes solution. WT and CD47^-/-^ mouse platelets were isolated as previously described (81). Washed platelets were resuspended in monocyte culture media and co-culture with monocytes in indicated ratios. For antibody mediated ligand/receptor blocking, platelets were treated with anti-CD47 antibody (40 μg/ml, Bio X Cell), anti-P-selectin antibody (40 μg/ml, Biolegend), anti-CD40L antibody (40 μg/ml, Fisher Scientific), Annexin V (2.5 μg/ml, Biolegend), RGDS peptide (250 μg/ml, Fisher Scientific) or anti-TGFβ antibody (40 μg/ml, Fisher Scientific) for 30 minutes before co-incubation with monocytes.

To obtain platelet releasates, washed platelets were incubated with thrombin (1 U/ml, Cayman Chemical) for 10 minutes. Thrombin was neutralized by hirudin (1 U/ml, Cayman Chemical). Platelets were pelleted by centrifugation, and the supernatant was collected to treat mouse monocytes in culture. Platelet conditioned media was obtained by culturing washed platelets in culture media overnight and collecting supernatant after centrifugation.

### Cell culture

Primary mouse BM monocytes were isolated from femurs and tibias. The BM was flushed with isolation buffer (1×PBS, 2% FBS, 1mM EDTA; Fisher Scientific) using a 26-gauge needle (BD Biosciences), and RBCs were lysed with ACK Lysis Buffer (Fisher Scientific). Cell suspension was passed through 100-μm cell strainer (Fisher Scientific), and monocytes were isolated using EasySep Mouse Monocyte Isolation Kit (Stemcell Technologies). Primary mouse circulating monocytes were collected by retro-orbital bleed into EDTA blood collection tubes and RBCs were lysed with ACK Lysis Buffer. Monocytes were isolated by the EasySep Mouse Monocyte Isolation Kit (Stemcell Technologies). Isolated monocytes were lysed for RNA extraction or cultured in RPMI media with 1× Glutamax (Fisher Scientific), 10% FBS, and 1% penicillin/streptomycin and treated with LPS (50 ng/ml) for 24 hours. Supernatants were collected for cytokine measurement by ELISA (DuoSet, R&D Systems). Primary mouse BM monocytes were co-cultured with mouse platelets in 1:20, 1:40, 1:80 ratio for 24 hours, platelets washed away and monocytes stimulated by LPS (10 ng/ml, Sigma-Aldrich), CpG (10 μg/ml, ODN1826, Invivogen) Pam2CSK9 (100 ng/ml, Cayman Chemical) or IL-1β (100 ng/ml, Fisher Scientific) for 24 hours. Mouse monocytes were also co-cultured with platelets for 24 hours, platelets washed away and monocytes left in culture plate for 2 days before LPS challenge (10 ng/ml). For experiments with protein blocking and inhibition, primary mouse monocytes were pretreated for 1hr with Pargyline (0.1 μM, 0.5 μM), SAHA (10 nM, 100nM), L4 (5nM), MTA (1 mM), anti-PSGL1 antibody (20 μg/ml, Bio X Cell), anti-CD18 antibody (20 μg/ml, Bio X Cell), anti-CD11b antibody (20 μg/ml, Biolegend), anti-SIRPα antibody (40 μg/ml, Biolegend), PTP Inhibitor (2 μM, 4 μM, SHP inhibitor, Cayman Chemical), 2-DG (0.2 mM, Millipore Sigma), MK2206 (2 μM, AKTi, Cayman Chemical) or Rapamycin (2 nM, mTORi, Cayman Chemical) for 30 mins prior to co-culture with platelets. Platelets were washed away 24 hours after co-incubation and monocytes were treated with LPS and IL-6 and TNFα release were determined though ELISA (DuoSet, R&D Systems). Transcripts for *Il6*, *Tnfa*, *Il1b* were analyzed by qPCR.

THP-1 cells (ATCC) were seeded in 24-well plates with RPMI media and 1× glutamax (Fisher Scientific), 10% FBS (Invitrogen) and 1% penicillin/streptomycin (Invitrogen). Cells were incubated with human platelets in 1:20, 1:40 and 1:80 ratio for 24 hours, platelets washed away and treated with LPS (5 μg/ml) for 24 hours. IL-6 and TNFα release were determined by ELISA (DuoSet, R&D Systems).

Mouse bone marrow derived macrophages (BMDMs) were generated as described(82). BMDMs were co-incubated with platelets in 1:30, 1:90 ratio for 24 hours, platelets washed away and challenged with LPS (10 ng/ml) for 24 hours. IL-6 and TNFα release were determined by ELISA (DuoSet, R&D Systems).

For transwell experiments, mouse primary monocytes or BMDMs were seeded at 2×10^5^/well into the lower chamber of a 24-well plate (Costar). Washed mouse platelets were cultured in the lower chamber directly in contact with the target cells or in the upper chamber separated from the target cells by 0.4 μm pore membrane, which allows diffusion of small molecules, but not platelets. Upper chamber was taken away after 24 hours and monocytes/BMDMs were treated with LPS for 24 hours. Culture supernatants were collected for measurement of cytokine production.

Human peripheral blood was obtained from confirmed sepsis patients in the University of Rochester Medical Center. PBMCs were isolated from whole blood using Ficoll-Paque Plus (VWR) density centrifugation. From PBMCs, primary human monocytes were isolated using EasySep Human Monocyte Isolation Kit (Stemcell Technologies). Monocytes were lysed and RNA extracted for RT-qPCR.

### Monocyte-platelet aggregate

BM monocytes were incubated with WT platelets, CD47^-/-^ platelets overnight and then monocytes were washed 3 times. Monocytes were stained with anti-CD41 Ab (Biolegend) and monocyte-platelet aggregates were determined by flow cytometry.

### Monocyte depletion

Mice were injected retro-orbitally with 40 mg/kg of control liposome or liposomal clodronate (L-alpha-Phosphatidylcholine, Encapsula Nano Sciences) at 24 and 4 h, respectively, before being challenged with LPS. Monocyte depletion was confirmed by flow cytometry analysis. Mouse peripheral blood was collected and subjected to RBC lysis. Cells were resuspended in PBS and incubated with APC-labelled anti-CD11b and PE-Cy7 labelled anti-Ly6C antibodies.

### qPCR

Primary mouse monocytes, circulating monocytes from sepsis patients were pelleted and resuspended in RLT lysis buffer. RNA was extracted using RNeasy Mini Kit (Qiagen). RNA concentration was measured using Nanodrop (Fisher Scientific). RNA was reverse transcribed into cDNA using High Capacity RNA-to-cDNA Kit (Applied Biosystems). Real-time PCR was performed using TaqMan Gene Expression Master Mix protocol on a Bio-Rad CFX Connect Real Time System. TaqMan Gene expression primers were purchased from Thermo Fisher Scientific.

### Proliferation assay

Primary mouse monocytes and BMDMs were seeded at 1×10^5^/well in 24-well plate. Cells were stained by 5μM CellTrace^TM^ CFSE (Fisher Scientific) for 20 minutes. Cells were washed twice with culture medium and co-cultured with washed platelets for 24 hours before LPS (50 ng/ml) stimulation. 24 hours after LPS treatment, cells were collected and proliferation was quantified by flow cytometry analysis. To study platelet mediated durable effect on monocytes, CFSE stained monocytes were left in culture for 2 days after platelet co-incubation. Then monocytes were treated with LPS (50 ng/ml) for 24 hours and cell proliferation was quantified by flow cytometry.

### Platelet transfusion

WT and CD47^-/-^ platelets were isolated as previously described. 1×10^9^ washed platelets were resuspended in 0.1 ml saline and injected retro-orbitally into each recipient mouse.

### RNA-Seq library preparation

Primary mouse monocytes were seeded at 2×10^5^ in 24-well plate. Monocytes were cultured alone or with 1×10^7^ washed mouse platelets for 24 hours. Platelets were washed away and monocytes were treated with PBS or LPS (50 ng/ml) for 4 hours. Total RNA from monocytes was extracted using the methods described above. To digest genomic DNA, mRNA was treated with DNase and processed for RNA-Seq with TruSeq Stranded mRNA library preparation by University of Rochester Genomic Research Core.

### Analysis of RNA-Seq data

R-4.0.2 was used for RNA-Seq data normalization and differential expression analysis. The heatmap was generated using pheatmap V1.0.12, using the significantly differentially expressed genes within the specific comparison. EnrichR GSE was performed on the significantly up or down regulated genes for a given comparison using EnrichR V3.0.

### Glycolytic rate assay

Primary mouse monocytes were seeded at 5×10^4^ in XFe96 microplates. Cells were co-incubated with control medium, 5×10^6^ mouse WT platelets, anti-CD47 antibody treated WT platelets or CD47^-/-^ platelets for 24 hours. Platelets were washed away before extracellular flux analysis was performed. The culture medium was switched to XF RPMI base medium (Agilent) supplemented with glucose (10 mM), pyruvate (1 mM), and glutamine (2 mM). Cells were incubated in a non-CO_2_ incubator at 37 °C for 1 hour. The extracellular acidification rate (ECAR) and oxygen consumption rate (OCR) were measured by XFe96 analyzer after sequential injection of the compounds from Seahorse XF Glycolytic Rate Assay Kit (Agilent) (rotenone and antimycin A (0.5 μM) and 2-DG (50 mM)). The ECAR and OCR were calculated by Seahorse Wave software (Agilent).

### Immunoblotting

Cells were lysed with 1×RIPA lysis buffer (EMD Millipore) mixed with protease inhibitor cocktail (MilliporeSigma). Cell lysis was centrifuged and supernatant was collected and mixed with Laemmli buffer (Bio-Rad). Protein samples were loaded into Mini-PROTEAN TGX Gels (Bio-Rad) and gels were run at 80 V, in 1×Tris/Glycine/SDS buffer (Bio-Rad). Protein was transferred from gel to PVDF membranes (Bio-Rad) in transfer buffer at 300 mA for 130 mins with ice packs. Blots were blocked in 5% BSA (MilliporeSigma) dissolved in Tris-buffered saline (Fisher Scientific) with 0.1% Tween-20 (TBST) for 1 hour at room temperature. Blots were incubated with primary antibody (1:1000, Cell signaling) overnight at 4 °C. Blots were washed and incubated in secondary anti-rabbit HRP antibody (1:2500, Proteintech) for 1 hour at room temperature. Immunoblots were developed with Supersignal West Pico (Fisher Scientific) using the Bio-Rad ChemiDoc MP chemiluminescence settings.

### ATAC-Seq library preparation

Circulating monocytes were isolated from 3 WT mice and 3 thrombocytopenic PF4-DTR mice using the methods described above. ATAC-Seq libraries were prepared from aliquots of 50,000 cells using the method described in Corces et al(83). Briefly, cells were incubated in hypotonic buffer in the presence of digitonin, washed, centrifuged to collect permeabilized samples, and resuspended in tagmentation buffer with Tn5 transposase (Illumina). Tagmented DNA was isolated with the Qiagen MinElute Reaction Cleanup kit and a dual-indexed library was prepared by subsequent PCR with Illumina-compatible primers and purified with AMPure XP beads (Beckman Coulter). Library quantity and quality were assessed by Qubit and Bioanalyzer (Agilent) assays, respectively. Paired-end sequencing was performed on an Illumina NovaSeq 6000.

### Analysis of ATAC-Seq data

Raw reads generated from the Illumina base calls were demultiplexed using Bcl2fastq version 2.19.1. Quality filtering and adapter removal was performed using FastP version 0.23.2(84). Processed/cleaned reads were then mapped to the mg38 + gencode M25 reference using bowtie2 version 2.4.4(85). Insert sizes were determined using Picard version 2.17.0. Deeptools version 3.1.3 aligmentSieve was used to filter for Encode blacklist regions (mm10) nucleosome free and nucleosome (mono, di, tri) regions(86). Broad peak enrichments were called with MACS2 using the following parameters “--broad -f BAMPE -g mm” for both nucleosome free and nucleosome regions(87). To determine differentially accessible regions, DiffBind and DESeq2 packages were used to compare broad peaks in sample groups(88, 89). Regions with an adjusted p-value < 0.05 were considered significant. The ChIPSeeker package was used to annotate genomic regions for each peaks(90). Custom plots were generated within the R software environment and the ggplot package.

### Data analysis

Flow cytometer was Accuri C6. All FACS data was analyzed using FlowJo version 7.6. ELISA results were measured by Plate Reader and ELISA data was analyzed by MyAssay. Real-time PCR results were analyzed in Microsoft Excel by calculating fold change 2^(-ΔΔCt)^ and *Actb* was used as a reference gene. Western blot data was quantified by ImageJ.

### Statistics

Statistics were computed in GraphPad Prism. Data were first subjected to a normality test; Standard 2-tailed Student’s *t* test was used when comparing 2 independent groups. One-way ANOVA followed by Bonferroni correction was used when comparing more than 2 independent groups. *P<0.05; **P<0.01; ***P<0.001. All data was represented as mean ± SEM.

### Study Approval

All human and animal studies were approved by the University of Rochester by our Institutional Research Board and Institutional Animal Care and Use Committee respectively. Written informed consent was received prior to participation of all human subjects.

## Author Contributions

CL designed studies, conducted experiments, acquired data, analyzed data, and contributed to writing of the manuscript; SKT, BN, SKBN, PM, ACL all contributed to conducting of experiments, MS performed human data statistical analysis, TJS performed genetic data acquisition and analysis, MK and APP acquired data and provided reagents, and CNM designed studies, analyzed data, provided reagents and contributed to writing of the manuscript.

## Acknowledgements

CNM has grants from NIH – R01HL160610, R01 HL141106 as well as Pilot Project Funding from the URMC Translational Immunology and Infectious Disease Institute to support these studies.

## Notes

### Competing Interest Statement

The authors have declared no competing interest.

