## Supplemental data for "Thrombocytopenia Independently Leads to Monocyte Immune Dysfunction"

### Supplemental Figure 1

A.

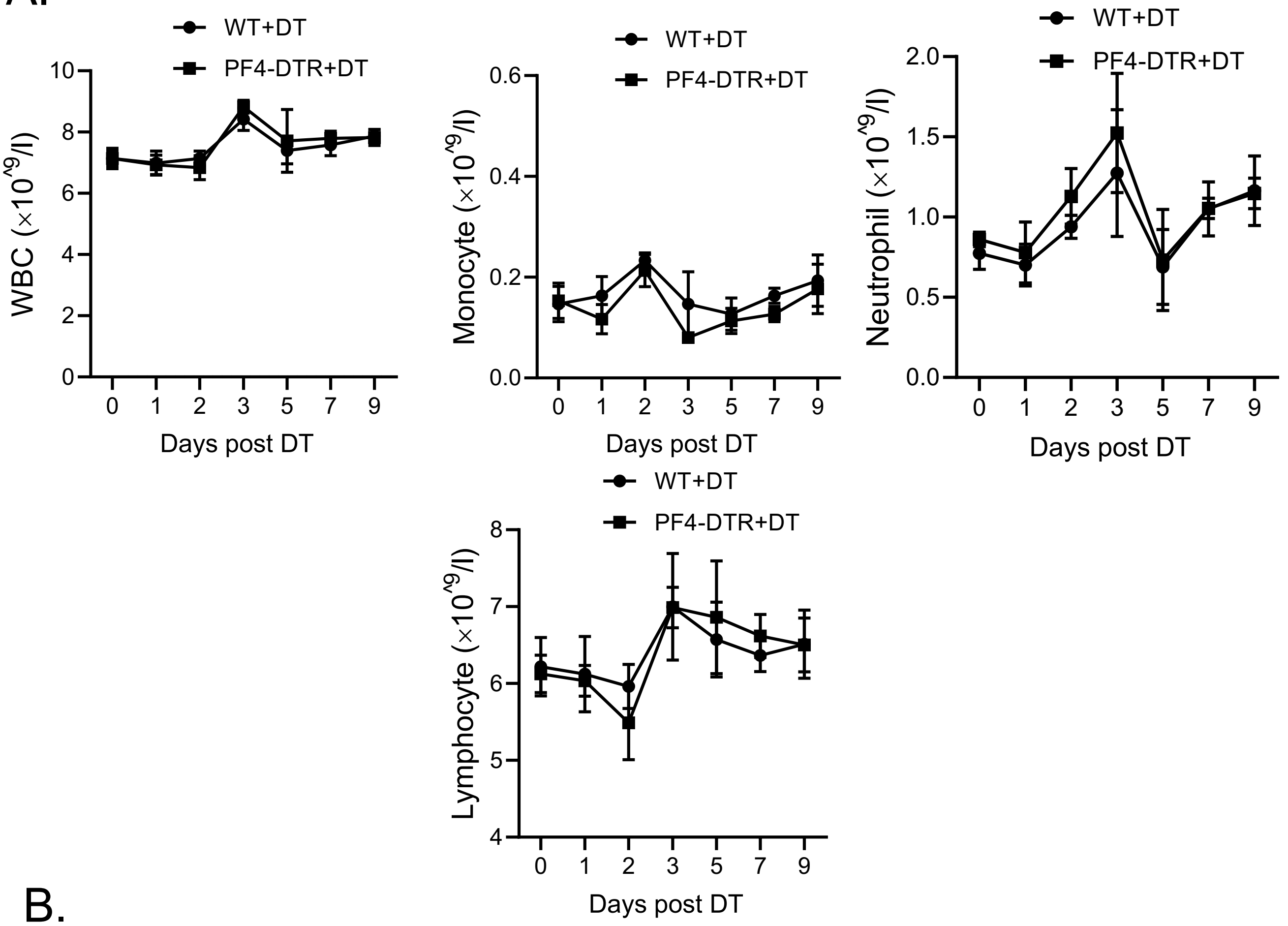


B.

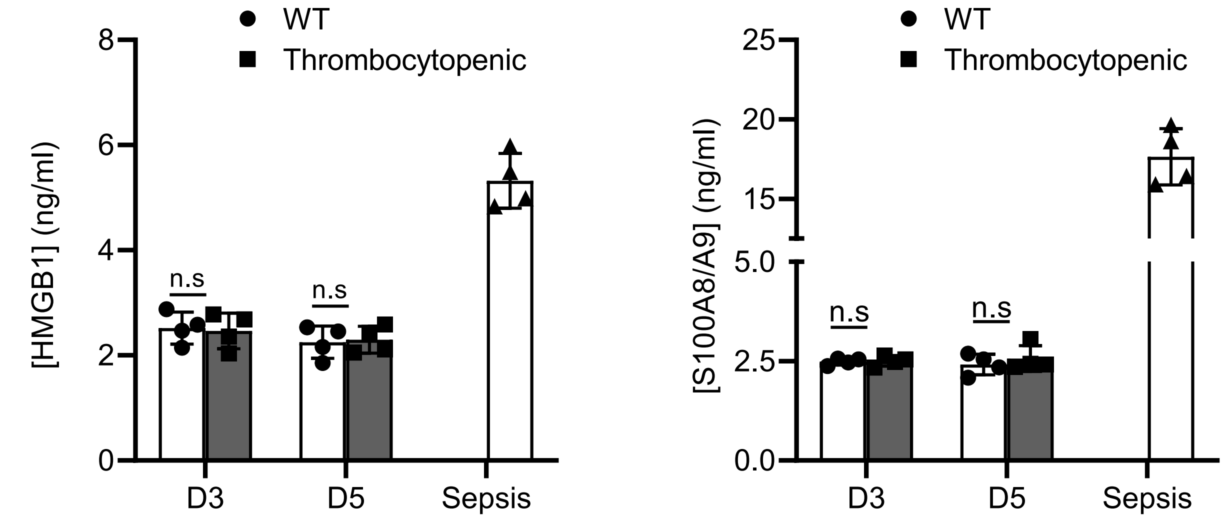

Supplemental Fig 1. A) Complete blood analysis on WT and PF4-DTR mice post DT. B) On d3 and d5 post-DT, WT and PF4-DTR mice had similar plasma levels of the DAMPS, HMGB1 and S100A8/A9 as determined by ELISA. Septic mouse plasma was used as a positive control. Data shown as mean ± SEM.

### Supplemental Figure 2

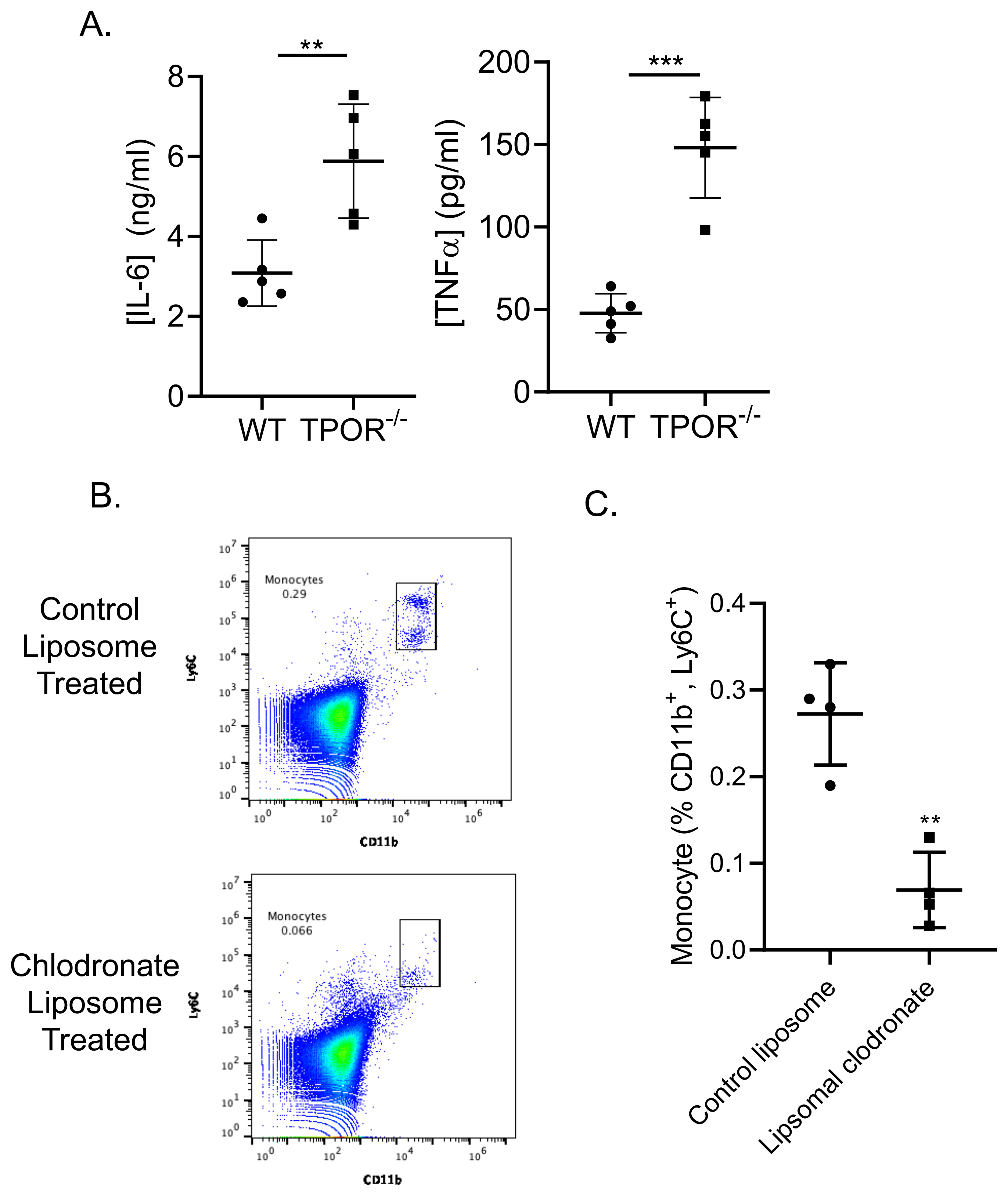

Supplemental Fig 2. A) $TPOR^{-/-}$ mice had an exaggerated response to polymicrobial sepsis challenge *in vivo*. Mice were treated with CS (0.8 mg/g) intraperitoneally and 4 hrs later plasma IL-6 and TNF$\alpha$ were determined by ELISA. (**P < 0.01, ***P<0.001, by unpaired, 2-tailed Student's *t* test). B-C) Liposomal clodronate depleted monocytes *in vivo*. Mice were injected retro-orbitally with control liposome or liposomal chlodronate (40 mg/kg) on d4 and d5 post-DT, and 4 hrs after the second chlodronate dose, blood was collected. B) Representative flow cytometry plots of circulating monocyte subsets and C) monocyte quantification. (**P < 0.01, by unpaired, 2-tailed Student's *t* test). Data shown as mean ± SEM.

### Supplemental Table 1

#### Sepsis patient characteristics[a]

| | |
|---|---|
| Age | 65 (58 – 70) |
| female sex | 7 (58) |
| Race | |
| Caucasian | 10 (84) |
| African- American | 1 (8) |
| Unknown | 1 (8) |
| Ethnicity | |
| Hispanic | 8 (67) |
| Non- Hispanic | -- |
| Unknown | 4 (33) |
| Charlson comorbidity index | 2 (1 – 3) |
| SOFA score, day 1 | 12<br>(10 – 14) |
| Source of infection | |
| Urinary tract | 5 (41) |
| Pulmonary | 3 (25) |
| Intra-abdominal | 2 (17) |
| Skin / soft tissue | 2 (17) |
| ICU length of stay (days) | 5.5 (3.9 – 13.3) |
| Hospital mortality | 2 (17) |

*[a]Values are median (interquartile range) or number (percent).*
*Definition of abbreviations: SOFA = sequential organ failure assessment ICU = intensive care unit.*

Table of confirmed sepsis patient characteristics.

### Supplemental Figure 3

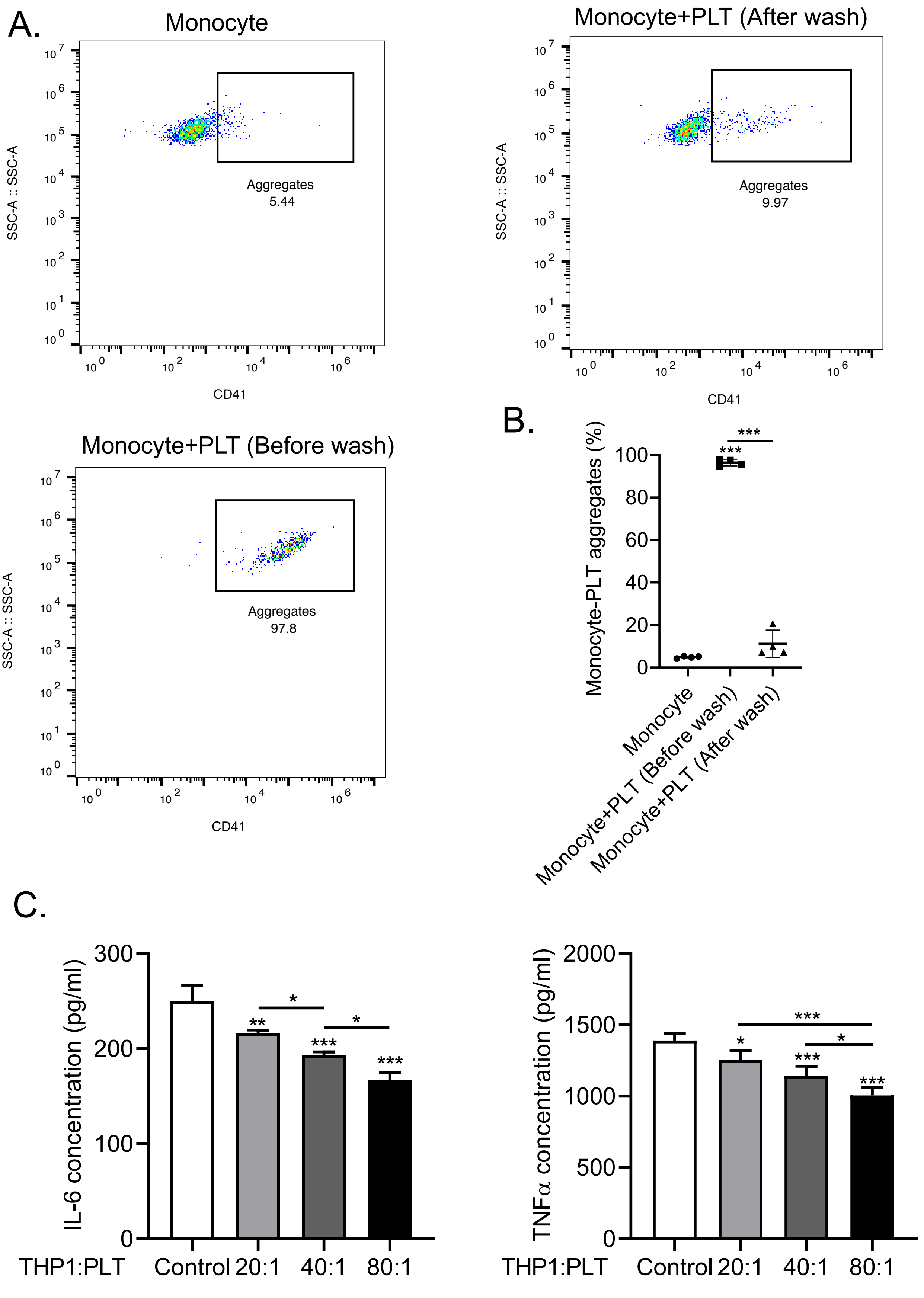

Supplemental Fig 3 A-B) Monocyte-platelet aggregates following co-culture and gentle washing. Mouse monocytes were co-incubated overnight with control media or platelets and the cells were then washed before monocyte-platelet aggregate quantification by flow cytometry. A) Representative flow cytometry plots and B) quantification. (***P < 0.001, by 1-way ANOVA followed by Tukey's post hoc test). C) Human monocyte cell line THP1 cells were co-incubated overnight with human platelets and cells were then washed to remove platelets. THP1 were then LPS stimulated and IL-6 and TNF$\alpha$ quantified. (*P < 0.05, **P < 0.01, ***P < 0.0001, by 1-way ANOVA followed by Tukey's post hoc test). Data shown as mean ± SEM.

### Supplemental Figure 4

A.

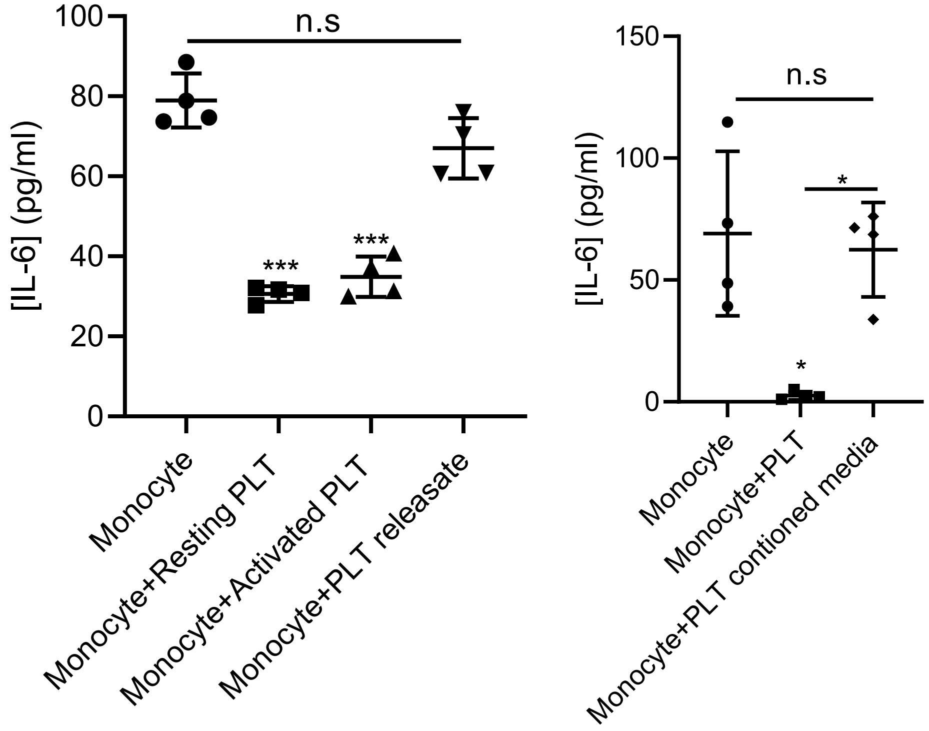


B.

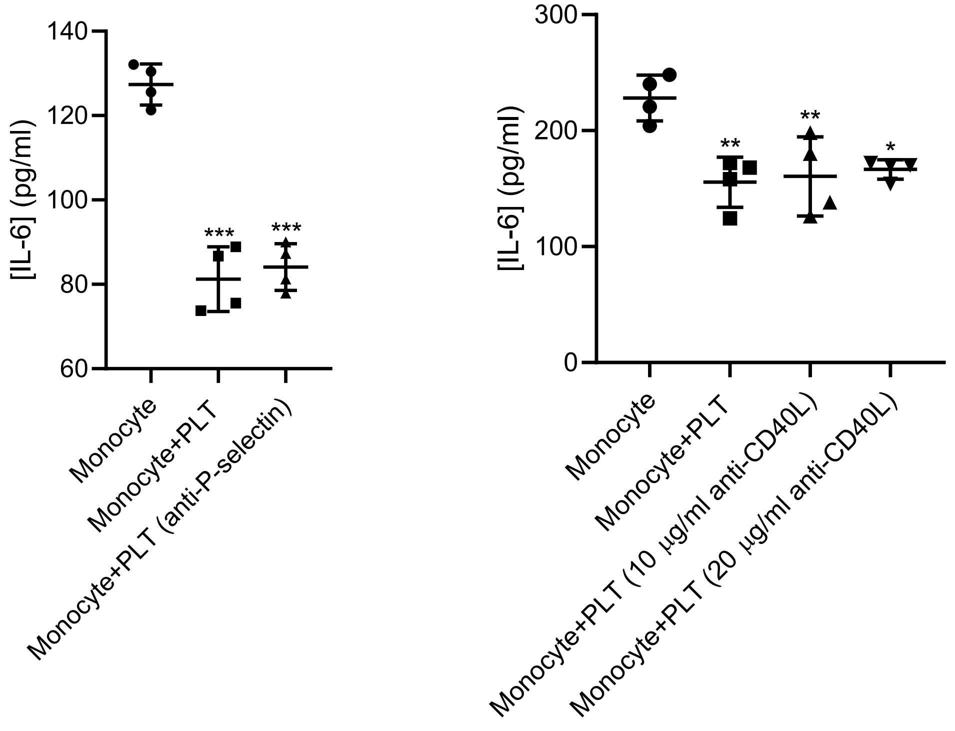

C.

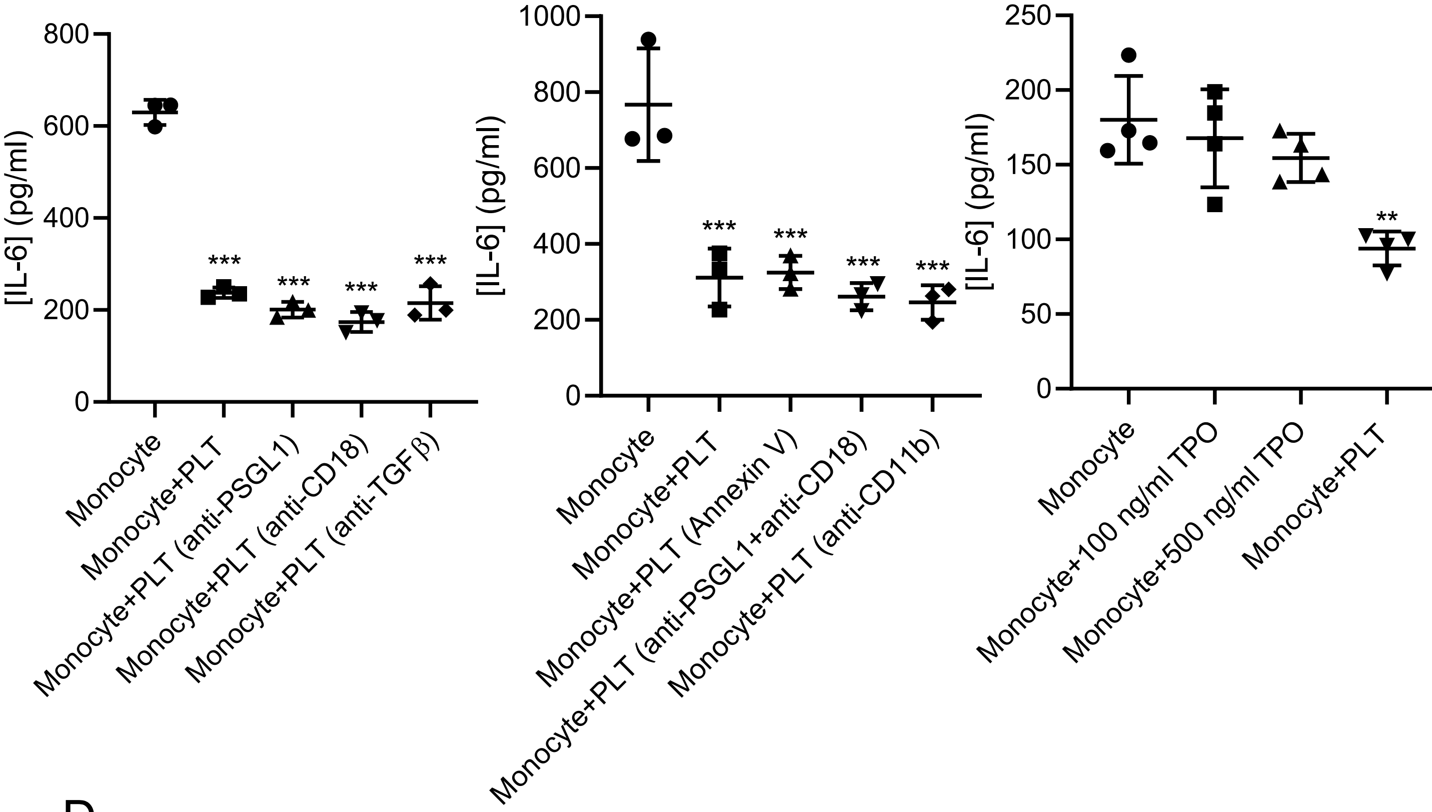

[IL-6] (pg/ml)
800
600
400
200
0
***
Monocyte
Monocyte+PLT
Monocyte+PLT (anti-PSGL1)
Monocyte+PLT (anti-CD18)
Monocyte+PLT (anti-TGFβ)
1000
Monocyte+PLT (Annexin V)
Monocyte+PLT (anti-PSGL1+anti-CD18)
Monocyte+PLT (anti-CD11b)
250
150
100
50
**
Monocyte+100 ng/ml TPO
Monocyte+500 ng/ml TPO


D.

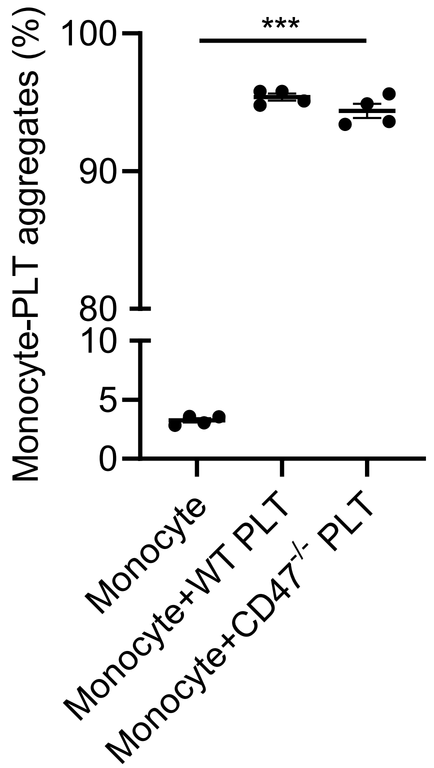

Monocyte-PLT aggregates (%)
100
90
80
10
5
0
***
Monocyte
Monocyte+WT PLT
Monocyte+CD47-/- PLT

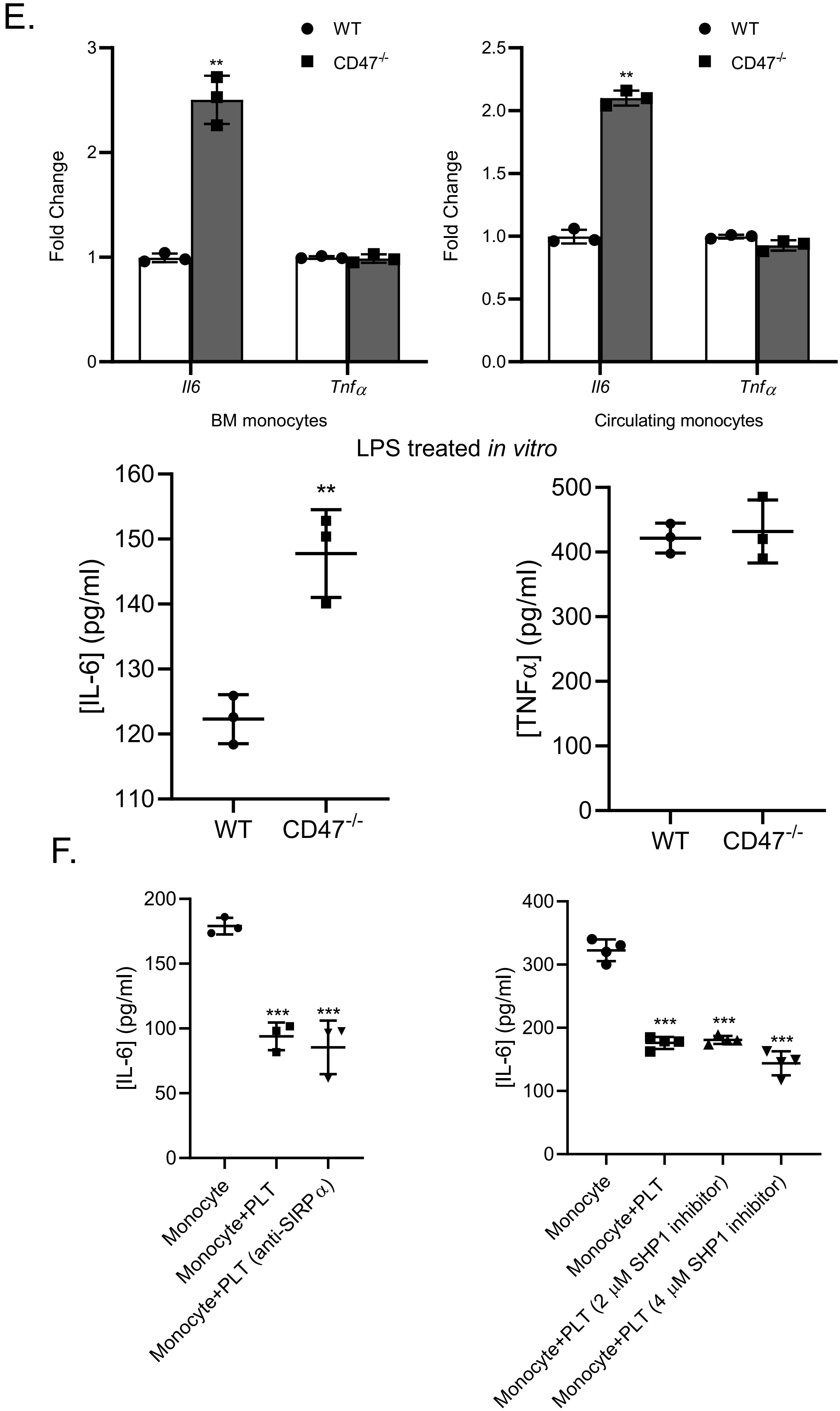

E.
WT
CD47-/-
Fold Change
Il6
Tnfα
BM monocytes
Circulating monocytes
**
LPS treated in vitro
[IL-6] (pg/ml)
[TNFα] (pg/ml)
WT
CD47-/-
F.
[IL-6] (pg/ml)
***
Monocyte
Monocyte+PLT
Monocyte+PLT (anti-SIRPα)
Monocyte+PLT (2 μM SHP1 inhibitor)
Monocyte+PLT (4 μM SHP1 inhibitor)

Supplemental Fig 4. Platelet mediators of monocyte immune tolerance. A) Monocytes were incubated with resting platelets, activated platelets, platelet releasates or platelet conditioned media overnight, platelets washed away and monocyte LPS stimulated. (*$P<0.05$, **$P<0.01$, ***$P<0.0001$, by 1-way ANOVA followed by Tukey's post hoc test). B) Monocytes were incubated with control platelets, P-selectin blocking Ab treated or CD40L blocking Ab treated platelets overnight and then LPS stimulated after platelets washed away. (*$P<0.05$, **$P<0.01$, ***$P<0.0001$, by 1-way ANOVA followed by Tukey's post hoc test). C) Monocyte were treated with PSGL-1 blocking Ab, CD18 blocking Ab or CD11b blocking Ab prior to platelet co-culture. After 24 hrs, platelets washed away and monocyte LPS stimulated. Monocyte were also treated with TPO or incubated with control platelets, TGF-β blocking Ab or Annexin V treated platelets overnight before LPS stimulation. (**$P<0.01$, ***$P<0.0001$, by 1-way ANOVA followed by Tukey's post hoc test). D) Monocytes form similar aggregates with $CD47^{-/-}$ platelets compared to WT platelets. Mouse monocytes were co-incubated overnight with WT or $CD47^{-/-}$ platelets and monocyte-platelet aggregates quantified by flow cytometry. E) Platelet mediated monocyte immune tolerance is CD47 dependent *in vivo*. Peripheral or BM monocytes were isolated from WT or $CD47^{-/-}$ mice for qRT-PCR, and circulating monocytes were stimulated with LPS *ex vivo*. (**$P < 0.01$, by unpaired, 2-tailed Student's *t* test). F) Blocking monocyte SIRP$\alpha$ signaling had no effect on platelet mediated monocyte LPS tolerance (*$P < 0.05$, **$P<0.01$, ***$P<0.0001$, by 1-way ANOVA followed by Tukey's post hoc test). Data shown as mean ± SEM.

### Supplemental Figure 5

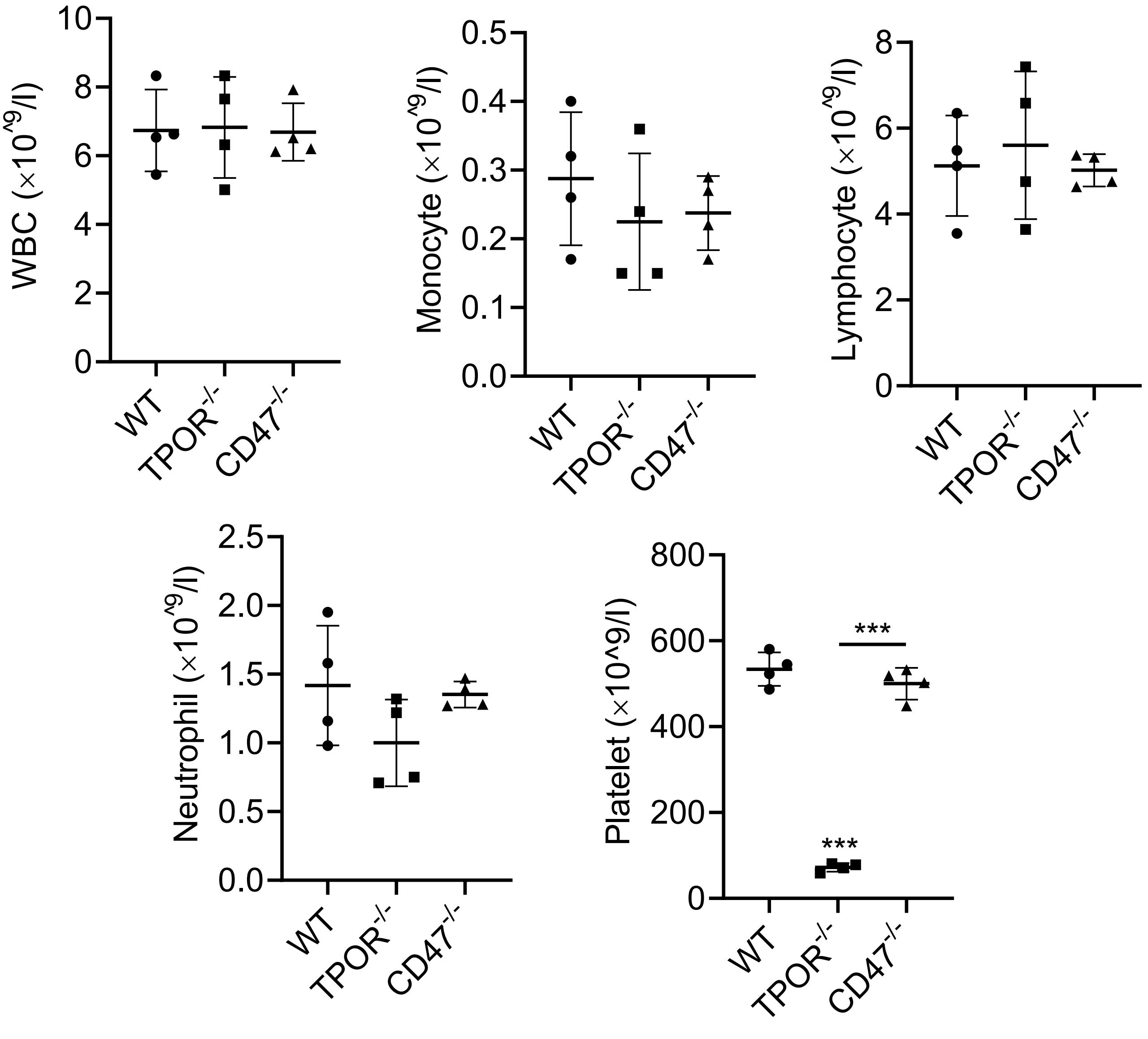


Supplemental Fig 5. Complete blood analysis on WT, $TPOR^{-/-}$ and $CD47^{-/-}$ mice.

### Supplemental Figure 6

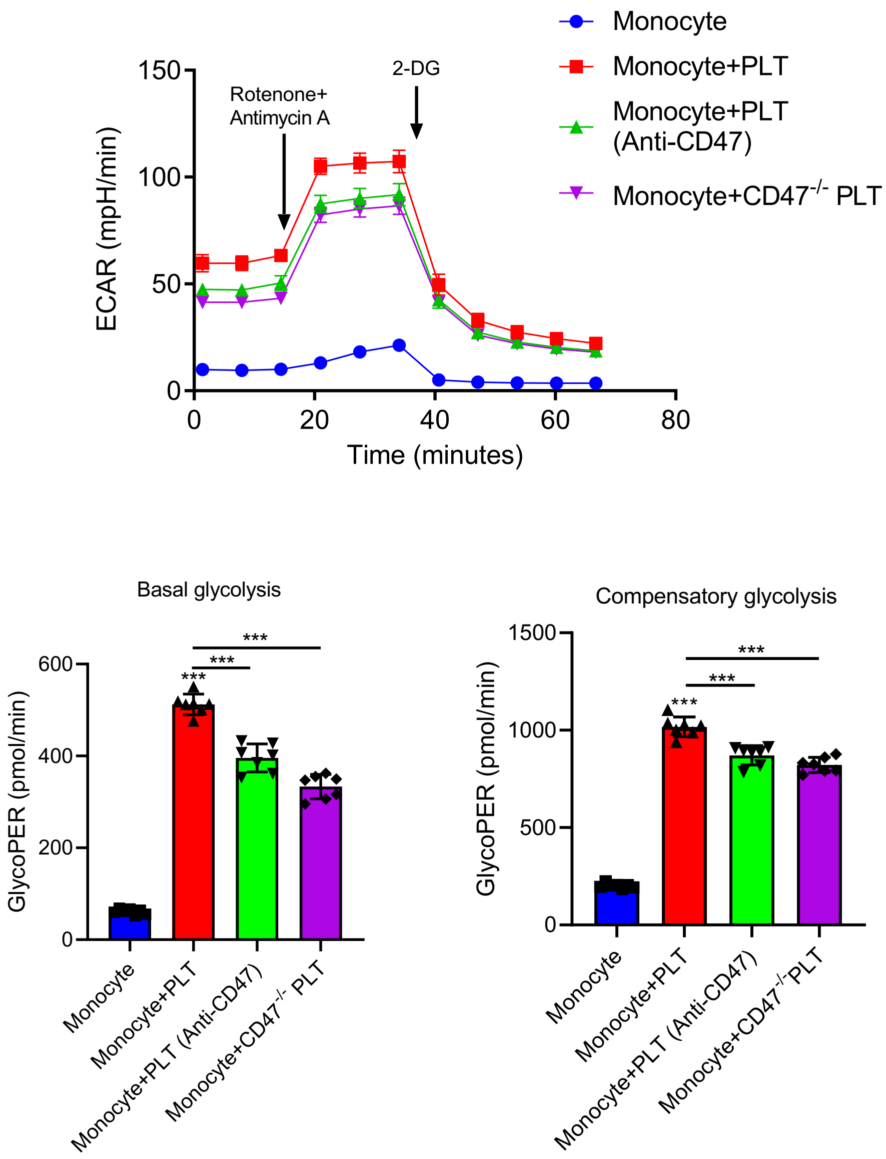


Supplemental Fig 6. Platelets increased monocyte basal and compensatory glycolysis in a CD47 dependent manner. Glycolytic rate assay was performed on monocyte incubated with control media, WT platelets, CD47 blocking Ab treated or CD47$^{-/-}$ platelets overnight. (***P < 0.0001, by 1-way ANOVA followed by Tukey's post hoc test).
